# Mosquito and primate ecology predict human risk of yellow fever virus spillover in Brazil

**DOI:** 10.1101/523704

**Authors:** Marissa L. Childs, Nicole Nova, Justine Colvin, Erin A. Mordecai

## Abstract

Many (re)emerging infectious diseases in humans arise from pathogen spillover from wildlife or livestock, and accurately predicting pathogen spillover is an important public health goal. In the Americas, yellow fever in humans primarily occurs following spillover from non-human primates via mosquitoes. Predicting yellow fever spillover can improve public health responses through vector control and mass vaccination. Here, we develop and test a mechanistic model of pathogen spillover to predict human risk for yellow fever in Brazil. This environmental risk model, based on the ecology of mosquito vectors and non-human primate hosts, distinguished municipality-months with yellow fever spillover from 2001 to 2016 with high accuracy (AUC = 0.72). Incorporating hypothesized cyclical dynamics of infected primates improved accuracy (AUC = 0.79). Using boosted regression trees to identify gaps in the mechanistic model, we found that important predictors include current and one-month lagged environmental risk, vaccine coverage, population density, temperature, and precipitation. More broadly, we show that for a widespread human viral pathogen, the ecological interactions between environment, vectors, reservoir hosts, and humans can predict spillover with surprising accuracy, suggesting the potential to improve preventative action to reduce yellow fever spillover and prevent onward epidemics in humans.

## Introduction

Many important (re)emerging infectious diseases in humans—including Ebola, sudden acute respiratory syndrome (SARS), influenza, *Plasmodium knowlesi* and other primate malarias, yellow fever, and leptospirosis—arise from spillover of pathogens from wildlife or livestock into human populations (1,2). While spillover is an important mechanism of human disease emergence, the drivers and dynamics of spillover are poorly understood and difficult to predict (3). Pathogen spillover requires favorable conditions to align in the reservoir (non-human animal), human, and pathogen populations and in the environment (3–5). Because these conditions interact, nonlinear relationships among the environment, host populations, and spillover probability are likely to emerge. Moreover, spillover is a probabilistic process that does not always occur, even when suitable conditions align. Despite these challenges, it is critical to predict pathogen spillover to enhance public health preparedness. Predicting spillover also provides an opportunity to test ecological approaches to solving globally important human health problems.

Most previous attempts to predict pathogen spillover have used statistical models (6–8). These models may be locally accurate for within-sample prediction, but may struggle to detect multidimensional, nonlinear, and stochastic relationships among host populations, pathogens, the environment, and spillover. In contrast, mechanistic models can test our understanding of transmission ecology, reproduce the complex, nonlinear interactions emerging in disease systems, and potentially improve our ability to predict spillover. In particular, Plowright *et al.* (3) recently proposed a mechanistic model, which remains untested, that integrates multiple ecological requirements to identify when conditions will align for pathogen spillover. Yellow fever in Brazil presents an ideal opportunity to test this model because the ecology of the pathogen has been studied for nearly 120 years (9), providing a wealth of mechanistic information and data, and because almost all recent cases in South America have occurred via spillover from the sylvatic cycle (10,11)

Yellow fever virus is a mosquito-borne *Flavivirus* that mainly persists in a sylvatic transmission cycle between forest mosquitoes (primarily *Haemagogus janthinomys, Hg. leucocelaenus,* and *Sabethes chloropterus* in South America) and non-human primates, and occasionally spills over into human populations (12). In some settings, these spillover events lead to onward human epidemics in an urban transmission cycle between humans and *Aedes aegypti* mosquitoes (9). Spillover of yellow fever requires the virus to be transmitted locally, mosquito vectors to acquire the virus from infected non-human vertebrate hosts, survive the extrinsic incubation period, and feed on human hosts, and human hosts to be susceptible to infection following exposure. These events require distributions of reservoirs, vectors, and humans, their interactions, and immune dynamics to align in space and time. In humans, yellow fever is the most severe vector-borne virus circulating in the Americas (10) with an estimated fatality rate for severe cases of 47% (13). While no urban transmission of yellow fever has occurred in the Americas since 1997 (14) and in Brazil since 1942 (15), a large epidemic began in December 2016 in Minas Gerais and by June 2018 had caused 2,154 confirmed cases and 745 deaths (16). Despite these large case numbers, molecular and epidemiological evidence suggests that human cases were caused by spillover from the sylvatic cycle, rather than urban transmission (11), most recently in areas previously believed to be free of yellow fever.

Prior statistical models have found climate and weather (including precipitation, temperature, and normalized difference vegetation index), non-human primate richness, land use intensiveness, and a latitudinal gradient to be predictive of the spatial and spatio-temporal distribution of yellow fever (6,8). We build on previous efforts by incorporating a mechanistic understanding of how ecological and human population factors affect yellow fever transmission and spillover. A mechanistic model allows for known relationships between the environment and transmission mechanisms, estimated from empirical data, to be included to test our understanding of the disease ecology. Additionally, mechanistic models allow for extrapolation beyond known regions to identify other regions where conditions are also suitable for yellow fever spillover. We use a mechanistic model encapsulating sylvatic yellow fever ecology to predict the spatial and temporal distribution of yellow fever spillover in Brazil, and we test the model on human yellow fever case data using a receiver operating characteristic curve and logistic regression. Here, we use “predict” to refer to independently estimating spillover risk mechanistically from simultaneous covariates and “forward prediction” to refer to estimating future spillover. We contrast this mechanistic prediction with statistical models that are fit to the spillover data, and therefore not able to make independent, out-of-sample predictions. We then incorporate the mechanistic model into further statistical analyses with boosted regression trees to understand what mechanisms our model does not capture.

Specifically, we ask: (1) Does the environmental suitability for sylvatic vectors, reservoir hosts, vector-human contact, and vector transmission—together termed environmental risk—predict geographic, seasonal, and interannual variation in yellow fever virus spillover into humans? (2) Are human population size and vaccine coverage, above and beyond environmental risk, critical for predicting spillover? (3) What additional environmental and population drivers might improve predictions of spillover? (4) Do the ecological processes that predict spillover in other parts of Brazil predict the recent yellow fever outbreak in the Southeast region of Brazil in 2016–2018, and if so, was risk elevated above historical baseline levels?

## Methods

Our goals were (1) to construct mechanistic estimates of yellow fever spillover risk over space and time, (2) to test these mechanistic risk models against observed cases of yellow fever spillover to humans, and (3) to statistically test for associations between observed spillover occurrence, mechanistically predicted risk, and environmental covariates to identify potential gaps in the mechanistic models. We constructed mechanistic risk estimates by modeling the ecological processes expected to drive transmission within reservoir hosts—vector distribution and seasonal abundance, vector dispersal, vector infectiousness, vector survival, vector–reservoir contact, and reservoir host distributions—and the risk of spillover to humans—human population density, vector–human contact rates, and human susceptibility (Fig. 1, Mechanistic model). For each of these ecological or human population factors, we parameterized a submodel using data from the literature and remotely-sensed covariates (Fig. 1 lists data sources and Fig. 2 shows the data and/or fitted submodels). We modeled several different risk metrics, as described below (see Methods: Spillover model). We then predict monthly risk of yellow fever spillover from the component submodels for each 1 km × 1 km pixel from December 2000 to December 2016 (Fig. 1; Supplementary Materials, Section 1.1). The risk estimates from January 2001 to December 2016 were aggregated to a municipality-level estimate to compare to available reports of human cases. Next, to test for relationships that were absent or mis-specified in our mechanistic model, we used both current and lagged aggregated municipality-wide environmental risk from December 2000 to December 2016 as covariates in a statistical model (a boosted regression tree) along with other environmental and demographic covariates to identify the traits of municipalities and months where yellow fever spillover occurred during the available human case data from 2001 to 2016 (Fig. 1, Statistical model). Finally, we sought to identify whether the mechanistic models predicted high suitability for spillover during the recent outbreaks (December 2016 – April 2018) (16). Given the limited time range of some covariates, we extrapolate model covariates for 2017 and 2018 by assuming that they were identical to 2016 or followed the same linear trend as was observed from 2015 to 2016. We then calculate the environmental risk metric for January 2017 to June 2018 in the region where the large outbreak occurred.

**Figure 1.**
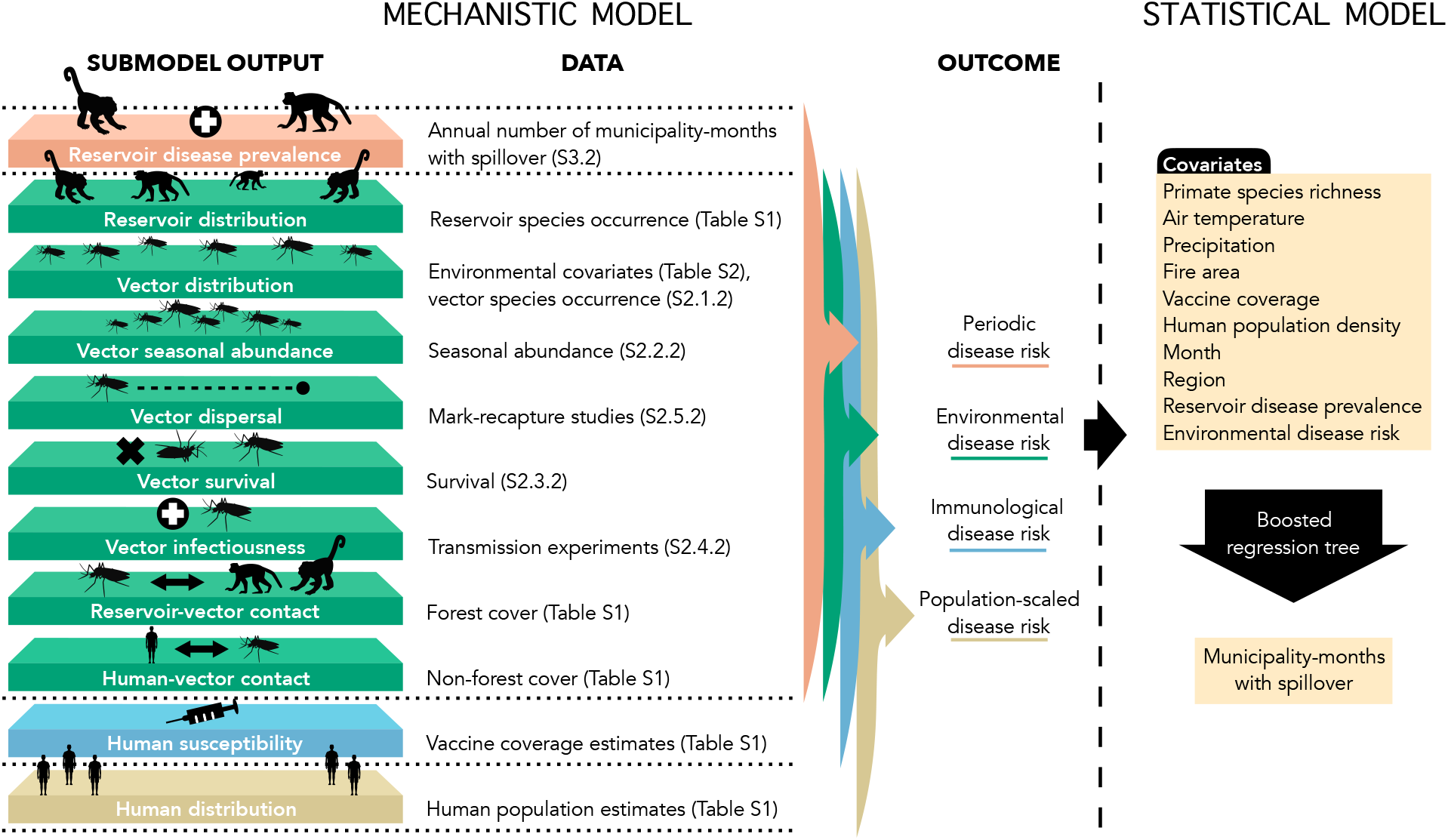
Mechanistic and statistical model schematic. Submodels of components in the mechanistic model are parameterized using independent data on reservoir species, vector species occurrences, seasonal abundances, vector mark-recapture studies, vector survival, transmission experiments, forest cover, estimated vaccine coverage, and human population estimates. Reservoir disease prevalence is estimated from annual number of municipality-months with spillover. The output from the submodels are used in a mechanistic spillover model to predict four metrics risk of yellow fever in humans: periodic disease risk, environmental disease risk, immunological disease risk, and population-scaled disease risk. Environmental disease risk metric is then used as a covariate in a boosted regression tree to predict the municipality-months with spillover and identify covariates important for predicting spillover. Other environmental covariates are also included in the boosted regression tree. Details on data used in the mechanistic model can be found in the Supplementary Materials (S). Specific locations within the Supplementary Materials are noted parenthetically by either the section or table in which details can be found. Data used in the boosted regression tree are described in the Supplementary Materials (Table S6). Layers shown on the left correspond to mechanistic model components in Fig. 2a–k.

### Spillover model

Yellow fever spillover risk is first estimated monthly from December 2000 to December 2016 using an adapted version of the model from Plowright *et al.* (3). We then estimate monthly spillover risk using extrapolated covariates (Supplementary Materials, Table S1) for the duration of the 2016 – 2018 outbreaks. We define environmental risk at a location 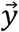 and time *t*—proportional to the number of infectious mosquito bites—as:

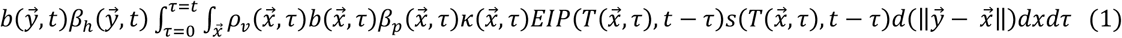

as a function of sylvatic vector density (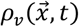, Fig. 2b and 2e), probability of biting non-human primates (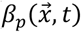, Fig. 2c) contingent on primate presence (Fig. 2a), probability of biting humans (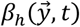, Fig. 2d) which depends on human presence (Fig. 2j), non-human primate infection prevalence 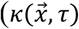, Fig. 2k), vector biting rate 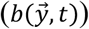, vector probability of becoming infectious 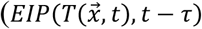, Fig. 2h), vector survival 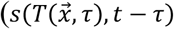, Fig. 2g), and vector dispersal 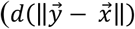, Fig. 2f), as described in Table 1. This model is a case study of a more general family of percolation models of pathogen spillover with alternative pathogen sources in space and time (Washburne *et al.,* this issue).

**Figure 2.**
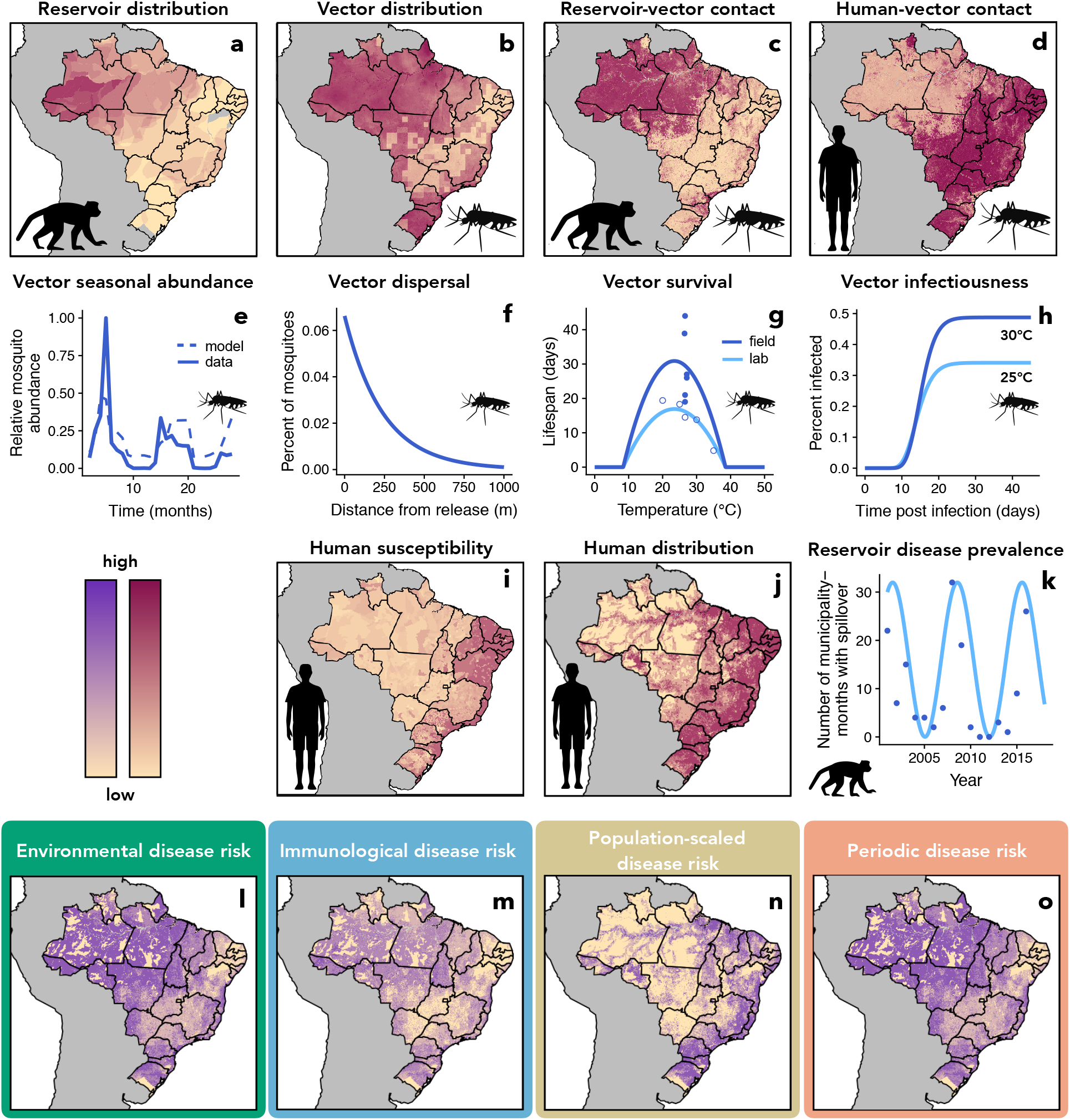
Data used to estimate ecological and human population components of spillover (a–k) and estimates of overall spillover risk (l–o). Number of primate reservoir species (a), vector species probability of occurrence (b), reservoir-vector contact probability (c), human-vector contact probability (d), human susceptibility approximated by one minus estimated vaccine coverage (i), and human distribution (j) vary spatially. Vector seasonal abundance is modeled as a function of rainfall using mosquito capture data (e). Vector dispersal depends on distance and is estimated from mark-recapture studies (f). Vector survival has been measured at different temperatures in laboratory (open circles) and field (closed circles) settings and was used to estimate temperature-dependent vector lifespan (g). Transmission studies at different temperatures inform modeled probability of vector infectiousness as a function of days since infecting bite and temperature (h). Phenomenologically modeled reservoir disease prevalence (light blue line, k) is approximated from human case data (blue dots, k). All mechanistic model components (a–k) are derived from empirical data in previously published studies. Components a–h are used to predict environmental risk of disease spillover (l), components a–i are used for immunological risk (m), components a–j are used for population-scaled risk (n) and components a–h and k are used for periodic risk (o). The four disease risk metrics presented here for illustrative purposes were estimated for January 2001 (l–o).

**Table 1.**
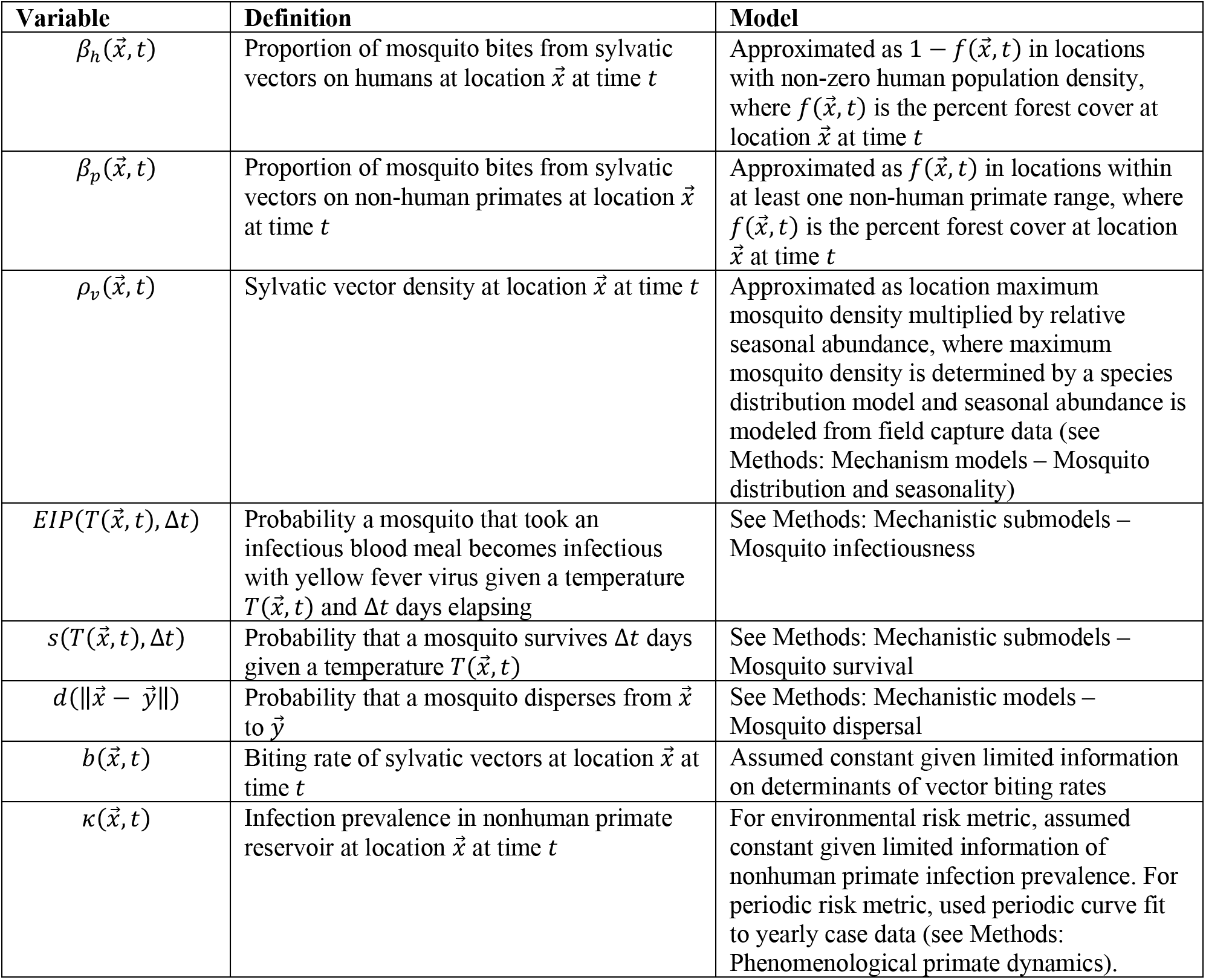
Spillover model variables and definitions.

We hypothesized that yellow fever spillover could be limited by environmental conditions, human susceptibility, human population distribution, and primate infection dynamics. To compare their relative importance, we define four metrics of model-predicted yellow fever spillover risk. First, we approximate *environmental risk* (Eq. 1, Fig. 2l), assuming that biting rate 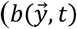 in Eq. 1,) and reservoir infection prevalence 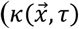 in Eq. 1) are constant over space and time in the absence of empirical data on these parameters, as described in Table 1. Since this metric ignores variation in human susceptibility, we then calculate *immunological risk* (Fig. 2m) as environmental risk multiplied by the estimated proportion of the human population that is susceptible to yellow fever (Fig. 2i), using previously estimated vaccine coverage rates (17). We then consider the influence of human population size on spillover risk by calculating *population-scaled risk* (Fig. 2n) as the immunological risk scaled by the number of people in a given location (Fig. 2j). Finally, we incorporate the effects of cycles of reservoir susceptibility and infection dynamics, for which data are not available, by calculating *periodic risk* (Fig. 2o), which uses a phenomenological periodic curve (Fig. 2k) for primate infection prevalence 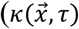 in Eq. 1). This periodic curve is designed to represent cycles of reservoir infection prevalence, driven by the demography of primate populations as naïve individuals are born, susceptible individuals accumulate, and epizootics become more likely (18). The full spillover model was run in Google Earth Engine (19). We estimate risk metrics monthly for 1 km × 1 km pixels using built-in functionality of Google Earth Engine that allows for calculations across differing scales by performing calculations for a specified output pixel scale.

### Mechanistic submodels

We fit mechanistic submodels from data for all key components of spillover (Fig. 1). For primate distribution (Fig. 2a), human susceptibility (Fig. 2i), and human population distribution (Fig. 2j), we used previously published estimates (17,20,21). All other mechanistic models (terms in Eq. 1) were fit with the R programming language, version 3.5.1 (22), with additional packages used for data processing, manipulation, and visualization (23–31).

Given limited information on the vector species, we use data for *Hg. janthinomys, Hg. leucocelaenus,* and *Sa. chloropterus* to fit models for the sylvatic vectors collectively for all mechanistic vector trait models. All data used were publicly available or results from previously published papers, as described in the Supplementary Materials (Table S1 and Mechanistic submodel details). Additional details on mechanistic model methods and data are available in the Supplementary Materials.

#### Vector distribution and seasonal density

To estimate the geographic distribution of sylvatic vector species (Fig. 2b), we fit a species distribution model (32,33) to *Hg. janthinomys, Hg. leucocelaenus,* and *Sa. chloropterus* occurrence data identified from the Global Biodiversity Information Facility (GBIF) (34,35) and a review of the literature (36–90), using the maxnet package in R (91). We included maximum, median, and minimum annual land surface temperature, total annual precipitation, precipitation in the driest month, precipitation in the wettest month, elevation, forest cover (%), land cover category, median annual enhanced vegetation index, and absolute latitude as predictors in the model (Supplementary Materials, Table S2). To account for uneven sampling effort across the geographic range, we corrected the background (pseudo-absence) points by subsampling from occurrence data of other mosquito species from GBIF (92). We calculated vector density as 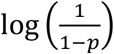, where *p* is the probability of occurrence estimated from the species distribution model (93). To estimate seasonal variation in vector abundance (Fig. 2e) due to rainfall seasonality (94), we fit a logistic regression of relative monthly vector abundance on current and one-month lagged relative monthly rainfall using field data (80,95–99) with glm in R.

#### Vector survival

To capture effects of temperature on vector survival (Fig. 2f), we used empirical data (100–102) and Bayesian inference to fit a quadratic function to the relationship between lifespan and temperature using rstan in R (103). Assuming constant vector mortality at a given temperature, we calculated daily survival probability as *p* = *e*^−1/*L*^, where *L* is vector lifespan (104).

#### Vector infectiousness

Virus infection, dissemination, and infectiousness in the vector are temperature-dependent (Fig. 2h) (105). We assume that vector competence—the probability that a vector exposed to an infectious blood meal becomes infectious with virus in its salivary glands—is a quadratic function of temperature, as shown for other flaviviruses (106). Additionally, we assume that at a given temperature, the extrinsic incubation period—the length of time required for an exposed vector to become infectious—is log-normally distributed across individuals (107,108). We fit a Bayesian model using experimental data (109–116) with the package rstan (103).

#### Vector dispersal

To estimate the range on which sylvatic mosquitoes disperse (Fig. 2f), we fit a negative binomial dispersal kernel (117) to mark-recapture data (118) using a Bayesian framework with the package rstan (103).

#### Vector contacts

We approximated reservoir-vector contact (Fig. 2c) as percent forest cover (119) contingent on the presence of at least one reservoir species (Fig. 2a). Similarly, we approximated human-vector contact (Fig. 2d) as percent non-forest cover (119) contingent presence of human population (Fig. 2j).

### Phenomenological primate dynamics

Primate population dynamics and susceptibility have been suggested as important constraints on yellow fever spillover (18), which remain poorly characterized. In the absence of primate infection data, we assumed that human spillover events are a proxy for infection prevalence during reservoir epizootics. This is the only mechanistic submodel that uses the human yellow fever spillover data directly—all other submodels are independent of human infection data. For this submodel, we used human cases of yellow fever reported by month of first symptoms and municipality of infection (2001–2016) from the Brazilian Ministry of Health (120). We define a spillover municipality-month as one in which at least one human case of yellow fever occurred. As an estimate of reservoir infection dynamics, we fit a phenomenological sine curve with a seven year period (121) to the yearly number of municipality-months with spillover (Fig. 2k) and then transformed the curve to be positive and less than one. The resulting curve is used as a spatially constant estimate of primate reservoir infection prevalence. Phenomenological primate dynamics are used in the periodic risk estimate (Fig. 2o) to account for a missing ecological process but are not used in any other risk metric, so all other risk metrics are parameterized independent of human spillover data.

### Model-data comparison

We compared spatially- and temporally-explicit mechanistic model predictions for spillover risk to observed human cases of yellow fever spillover using a statistical model. We limit the comparison to 2001–2016 based on the availability of human case data. We considered four modeled risk metrics (defined above): environmental risk, immunological risk, population-scaled risk, and periodic risk. Because risk was modeled by pixel, to compare the model output with municipality-month observations of human cases, we calculate both mean risk and maximum risk in each municipality and month. While mean risk may be more representative of the entire municipality, we hypothesized that maximum risk in the municipality-month might better predict the small-scale processes that drive spillover. The use of maximum risk also may also help to avoid spatial aggregation that can lead to bias or mask the relationships, for example the modifiable areal unit problem (122).

We compared municipality means and maxima for all four risk metrics to human yellow fever data for model evaluation in the following three ways. First, for each modeled risk metric and each municipality summary statistic (mean and maximum), we fit a logistic regression of spillover probability as a function of model-predicted risk (Supplementary Materials, Table S4) using glm in R (22). Second, we calculated a receiver operating characteristic curve to calculate the area under the curve (AUC), a measure of goodness of fit, for each modeled risk metric and municipality summary statistic (Supplementary Materials, Table S4). As this analysis focuses on prediction of spillover as a way to compare hypothesized mechanisms, comparison of AUC values to a null model is beyond the scope of this paper. Finally, for all eight mechanistic predictions and estimated vaccine coverage, we regressed the number of reported yellow fever cases given that spillover occurred and calculated Spearman’s rank correlation coefficient with number of reported cases to consider nonlinear but monotonic associations (Supplementary Materials, Table S5).

### Statistical model

We used a boosted regression tree (123,124) to understand any potential gaps in the mechanistic model and its relationship to environmental and human population covariates. As predictors of yellow fever spillover in the boosted regression tree, we included the following covariates for each municipality-month: current and one-month lagged maximum predicted environmental risk, current and one-month lagged fire area, average and maximum number of primate species, estimated municipality vaccine coverage, average human population density, average monthly air temperature, average monthly precipitation, phenomenological primate dynamics, region, and month (Supplementary Materials, Table S6). Each observation is a municipality-month and the response variable is the binary indicator of whether or not yellow fever spillover occurred in a municipality-month (see Methods: Model-data comparison). While some of predictor covariates contribute to the environmental risk metric (i.e., air temperature, rainfall, and primate reservoir ranges), we also include them in the boosted regression tree analysis to identify whether the environmental covariates have any predictive power beyond their role in the mechanistic model, which could indicate that the mechanistic model does not fully capture their influence on spillover. We included fire area as a proxy for land conversion (125), which has previously been shown to be predictive of yellow fever spillover (8). We also included vaccine coverage and human population density despite their poor predictive performance in the mechanistic model to identify whether these human population factors are predictive of spillover in ways not previously hypothesized, and therefore not captured in the mechanistic model. Boosted regression trees repeatedly fit regression trees, which create multiple binary splits in the dataset based on predictor variables. Each successive tree is fit to the residuals of the previous best model. The model is then updated to include the next tree (123). Variable importance is calculated as a weighted sum of the number of times a variable is used for splitting, with weights determined by the squared improvement due to the split (123).

We fit the boosted regression tree to data from 2001 to 2016, given this is the range of the available human case data for inferring spillover. We partition the dataset into spatially- and temporally-balanced training (80%) and test (20%) sets prior to the analysis. Optimal learning rate, tree complexity, and number of trees were selected as the set of parameters that minimized cross-validation predictive deviance (Supplementary Materials, Table S7; 63). The dataset was split in R using the BalancedSampling package (126), models were fit in R using the gbm and dismo (127,128) packages, and variable effects were calculated with the pdp package (129). Additional details can be found in the Supplementary Materials.

## Results

Primate species distribution (Fig. 2a), vector distribution (Fig. 2b; Supplementary Materials, Fig. S1 and S2), reservoir-vector contact (Fig. 2c), human-vector contact (Fig. 2d), and human susceptibility (Fig. 2i) varied over space and time based on estimates and models fit to empirical data. In addition, vector survival (Fig. 2g) and infectiousness (Fig. 2h, Supplementary Materials, Fig. S4 and S5) varied with temperature, vector abundance varied seasonally with rainfall (Fig. 2e; Supplementary Materials, Fig. S3 and Table S3), and vector dispersal declined exponentially with distance (Fig. 2f; Supplementary Materials, Fig. S6). Together, these empirical relationships between environment and host, vector, and virus ecology compose an estimate of environmental risk of yellow fever spillover (Supplementary File 2).

The environmental risk model strongly predicted episodes of yellow fever spillover into humans (AUC = 0.72) and adding phenomenological reservoir infection dynamics in periodic risk further improved the model (AUC = 0.79; Fig. 3). Surprisingly, models that included human vaccination coverage and human population size performed worse than the environment-driven models (AUC = 0.64 and 0.64; Fig. 3). For all risk metrics, maximum value in the municipality-month was a better predictor of spillover than mean value (Fig. 3). Logistic regressions of spillover probability as a function of model-predicted risk showed similar patterns in Akaike Information Criterion (AIC) values (Supplementary Materials, Table S4). Model-predicted environmental (mean and maximum), periodic (mean and maximum), and immunological (maximum) risk metrics were statistically significant predictors of spillover probability at the 5% level after correcting for multiple hypothesis testing (Supplementary Materials, Table S4; 71). By contrast, given that spillover occurred, none of the eight mechanistic model risk summaries were statistically significant predictors of number of cases, nor was estimated vaccine coverage (Supplementary Materials, Table S5). In Spearman’s rank correlations, we find that of the independent predictors, vaccine coverage is most correlated with risk, followed by maximum environmental risk, although these correlations are weak (Supplementary Materials, Table S5).

**Figure 3.**
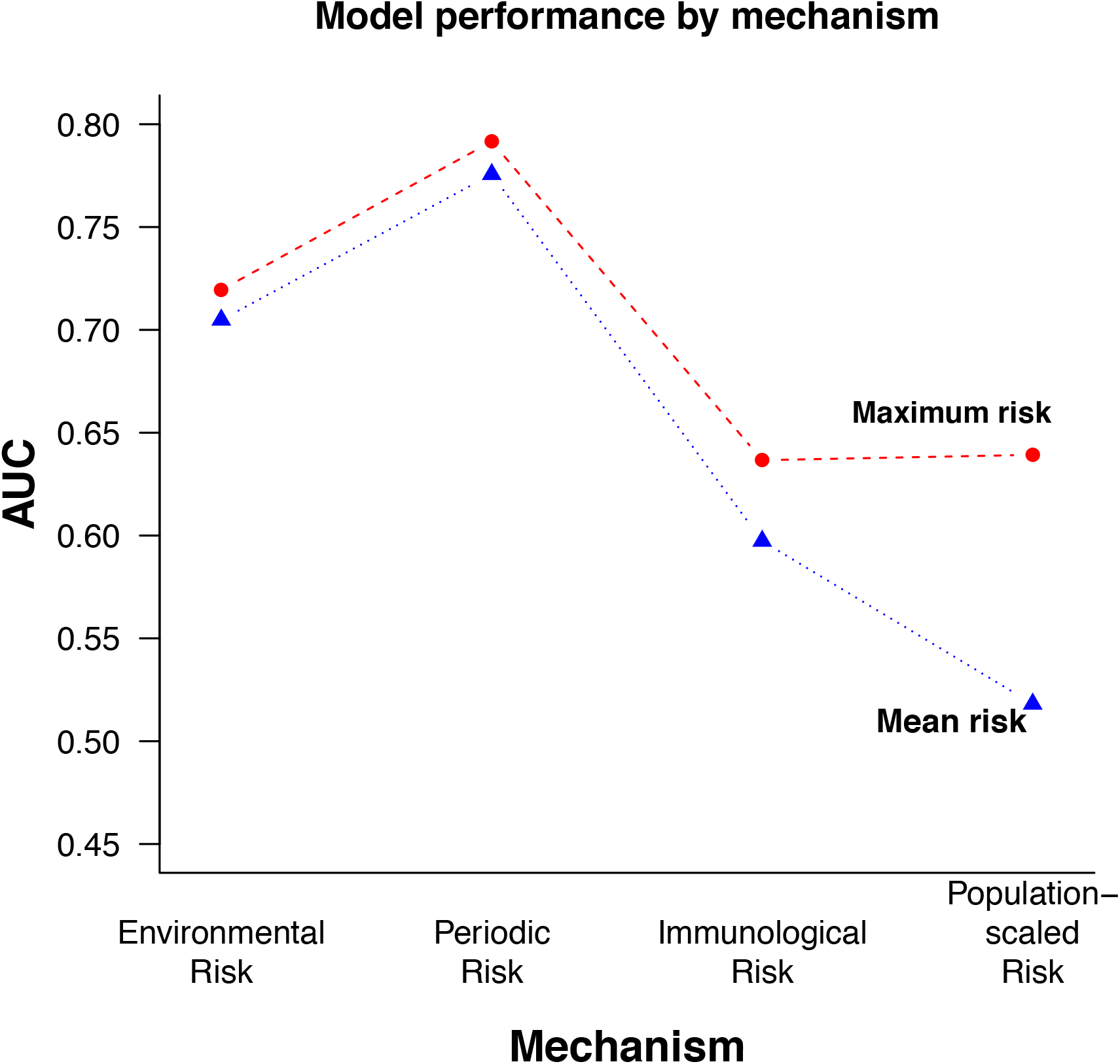
Municipality maximum periodic risk best predicts spillover. Each point is the calculated area under the curve (AUC) from spillover predicted by modeled risk, where higher AUC represents a model better able to distinguish between spillover and non-spillover observations. The risk models (from left to right on the *x*-axis) are environmental risk, periodic risk, immunological risk, and population-scaled risk. Municipality-wide maxima (red dashed lines and circles) and means (blue dotted lines and triangles) are shown for each metric.

Mechanistic model estimates matched seasonal variation in spillover, and accurately captured differences in seasonality by region (Fig. 4a). Risk peaked in April in the North and Northeast regions and in February in Central-West, South, and Southeast regions. The seasonal regional correlation between number of municipality-months with spillover and average environmental risk was highest in the Southeast (0.77), followed by the South (0.61), Central West (0.58), and North (0.42) regions. The periodic risk matched interannual variation in spillover (Fig. 4c), an unsurprising finding given periodic risk incorporated phenomenological primate dynamics derived from human cases of spillover. Interannual regional correlations were weaker than seasonal correlations but similarly highest in the Southeast (0.54), followed by the Central-West (0.45), North (0.21), and South (0.13) regions.

**Figure 4.**
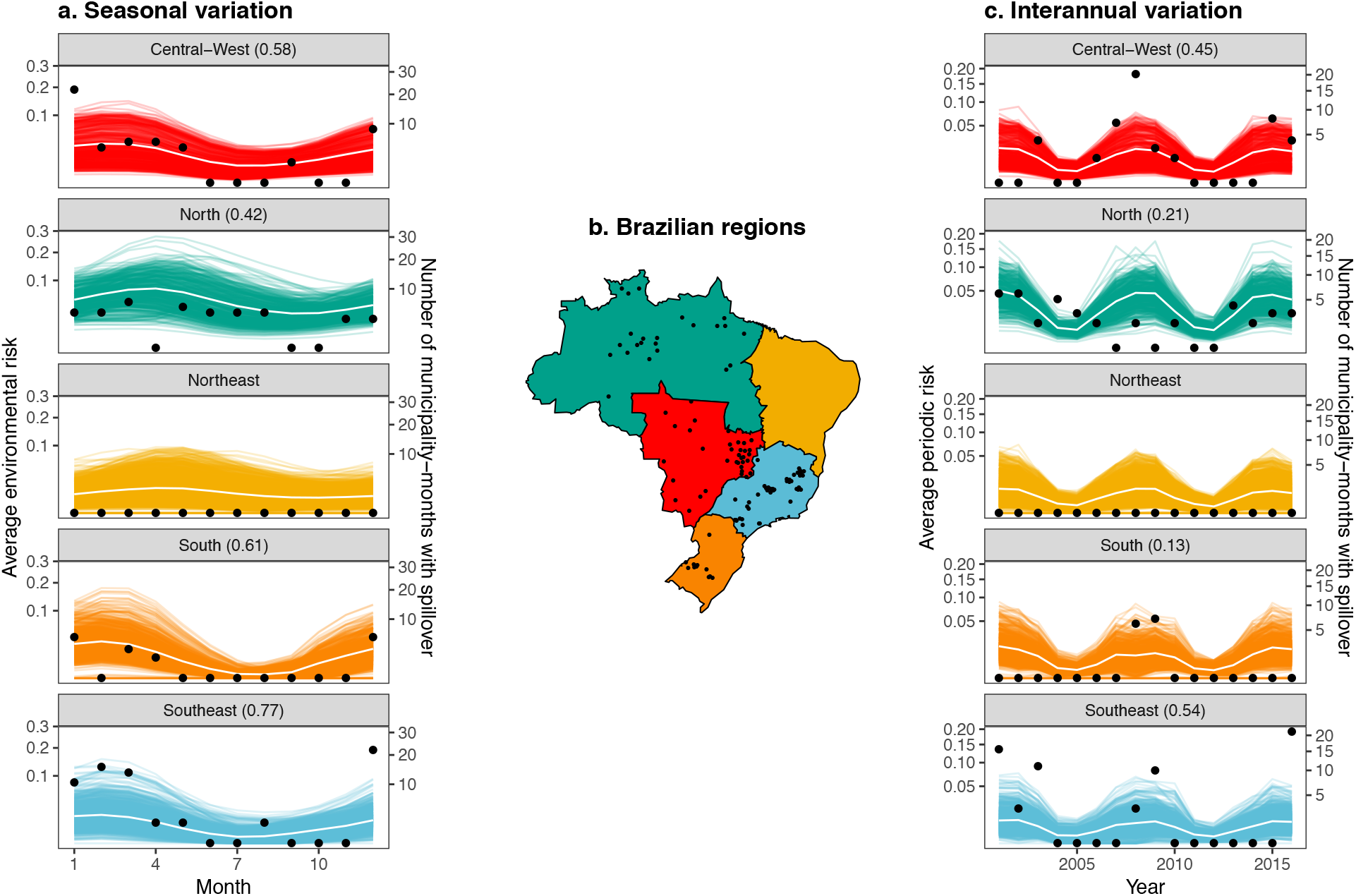
Modeled environmental risk captures seasonal variation and periodic risk captures interannual variation in spillover. Each colored line is the seasonal average of modeled maximum environmental risk in a municipality (a) and the yearly average of modeled maximum periodic risk in a municipality (c). White lines are the regional average over the municipal curves. Black points represent the total number of municipality-months with spillover in that region per month (a) and per year (c), or the municipalities with at least one month with spillover (b). Correlations between regional average environmental risk (white lines) and regional number of municipality-months with spillover (black points) shown in parentheses (a, b) for regions where spillover has occurred (all except the Northeast). Regions of Brazil are shown with corresponding colors (c). The Southeast (shown in blue) was the region with the majority of cases during the large outbreaks in 2016–2018.

The boosted regression tree found one-month lagged environmental risk and current environmental risk to be the second and fifth most important predictors of spillover, respectively (Fig. 5). Not surprisingly, the boosted regression tree significantly improved predictive performance from the mechanistic model because it was trained on the human spillover data (training AUC > 0.99, test AUC = 0.95). Vaccine coverage, temperature, population density and precipitation were also among the six most important predictors in the boosted regression tree. As expected, municipality-months with spillover had higher current and one-month lagged environmental risk (Fig. 5b and e), as well as high (phenomenologically) estimated primate infection prevalence and high primate species richness (Supplementary Materials, Fig. S8). We find that municipality-months with spillover have low monthly average temperatures (Fig. 5a), which may in part be due to poorly captured effects of temperature in the mechanistic model from averaging temperature before calculating mosquito trait values for survival and infectiousness (131). We also find that municipality-months with spillover have low rates of precipitation (Fig. 5d), which may correspond to settings with increased human activity in the forest, and therefore increased chance of spillover. However, current and lagged fire area, hypothesized indicators of deforestation activity, were not significant predictors of spillover in the boosted regression tree models (Supplementary Materials, Fig. S8).

**Figure 5.**
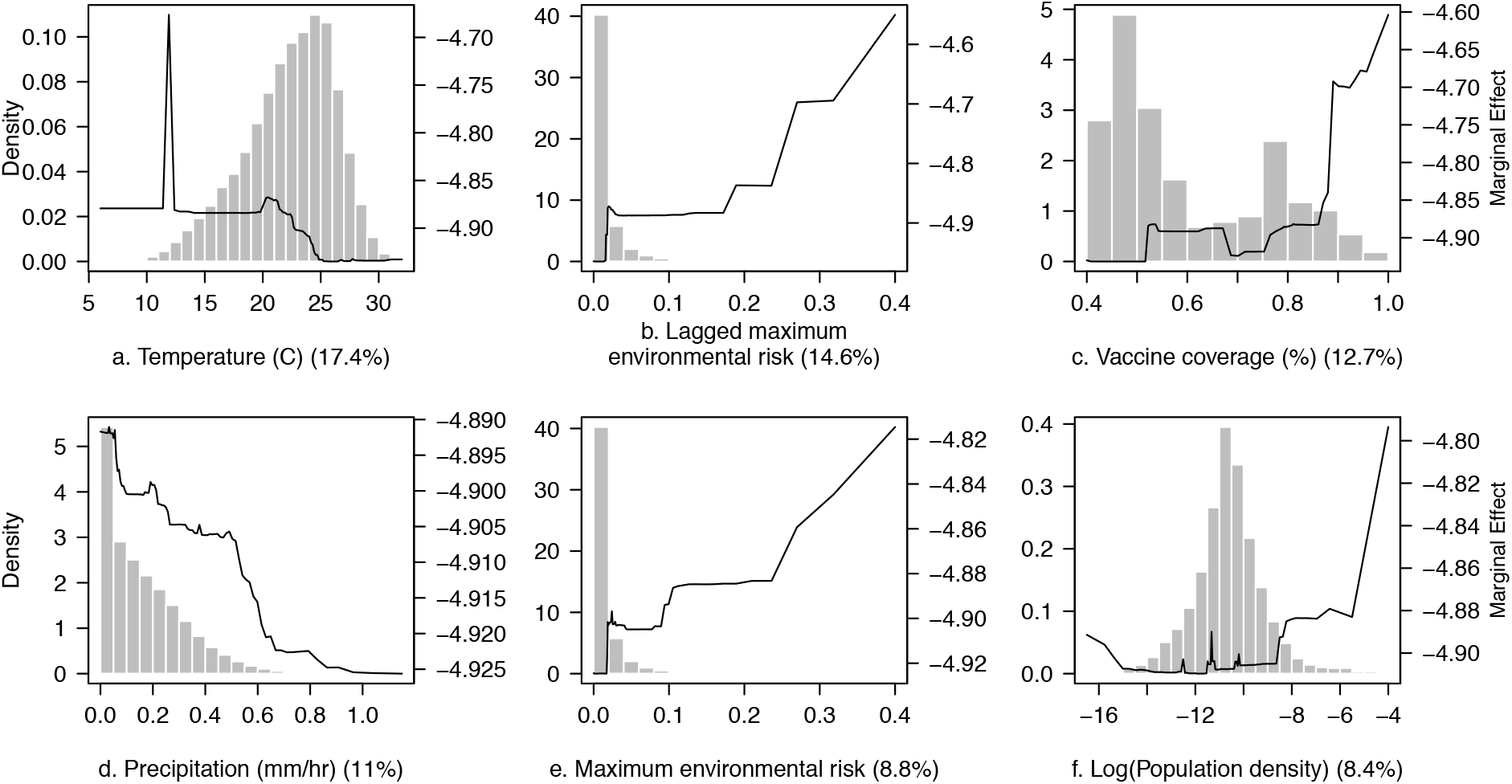
Partial dependence plots of top six predictors of spillover in a municipality-month from boosted regression tree analysis. Plots are listed in order of predictive importance with relative influence (%) listed. In order, the variables identified as most important predictors were average temperature in the municipality-month (a), one-month lagged maximum environmental risk (b), estimated vaccine coverage (c), average rate of precipitation in the municipality-month (d), current month maximum environmental risk (e), and municipality population density (log-scaled for visibility, f). Histograms show the distribution of observed municipality-months at each covariate value (left *y*-axis) and solid lines show the marginal effects of covariate on model prediction (right *y*-axis). Marginal effects highlight the characteristics of municipality-months that experienced spillover in Brazil 2001–2016.

Unexpectedly, municipality-months with spillover tended to have vaccine coverage above 90%, suggesting that high rates of vaccine coverage do not prevent spillover from occurring. While estimated vaccine coverage was included as a measure of human susceptibility, it is likely capturing other patterns in the spatial distribution of spillover; regions known to experience yellow fever spillover are likely to have high vaccination rates, while those where spillover is rare or nonexistent are likely to have low vaccination rates. Accordingly, estimated vaccine coverage is bimodal, potentially due to a group of lower risk municipalities and a group of higher risk municipalities. The partial dependence plot also displays two plateaus in the marginal effect in the vaccine coverage on model estimates, which roughly correspond to the two vaccine coverage groups.

The recent outbreaks in Brazil in the 2016–2017 and 2017–2018 transmission seasons have been the largest in over 50 years (16). The environmental risk model predicts persistent, low environmental risk of spillover in the affected states (Minas Gerais, Espírito Santo, Sao Paulo, and Rio de Janeiro) and does not predict any increase in spillover risk during the recent transmission seasons (Fig. 6). The date ranges of confirmed human cases during the 2016–2017 and 2017–2018 outbreaks are shown in pink bands (Fig. 6) based on a World Health Organization epidemiological update (16). The mechanistic model predicts spillover risk in Espírito Santo and Rio de Janeiro, where no spillover occurred from 2001 to 2016, at levels similar to those in Minas Gerais and Sao Paulo, where spillover had previously occurred. As in other regions, the model accurately captures the seasonality of spillover risk in this region (Fig. 6), which is distinct from that of other regions (Fig. 4a).

**Figure 6.**
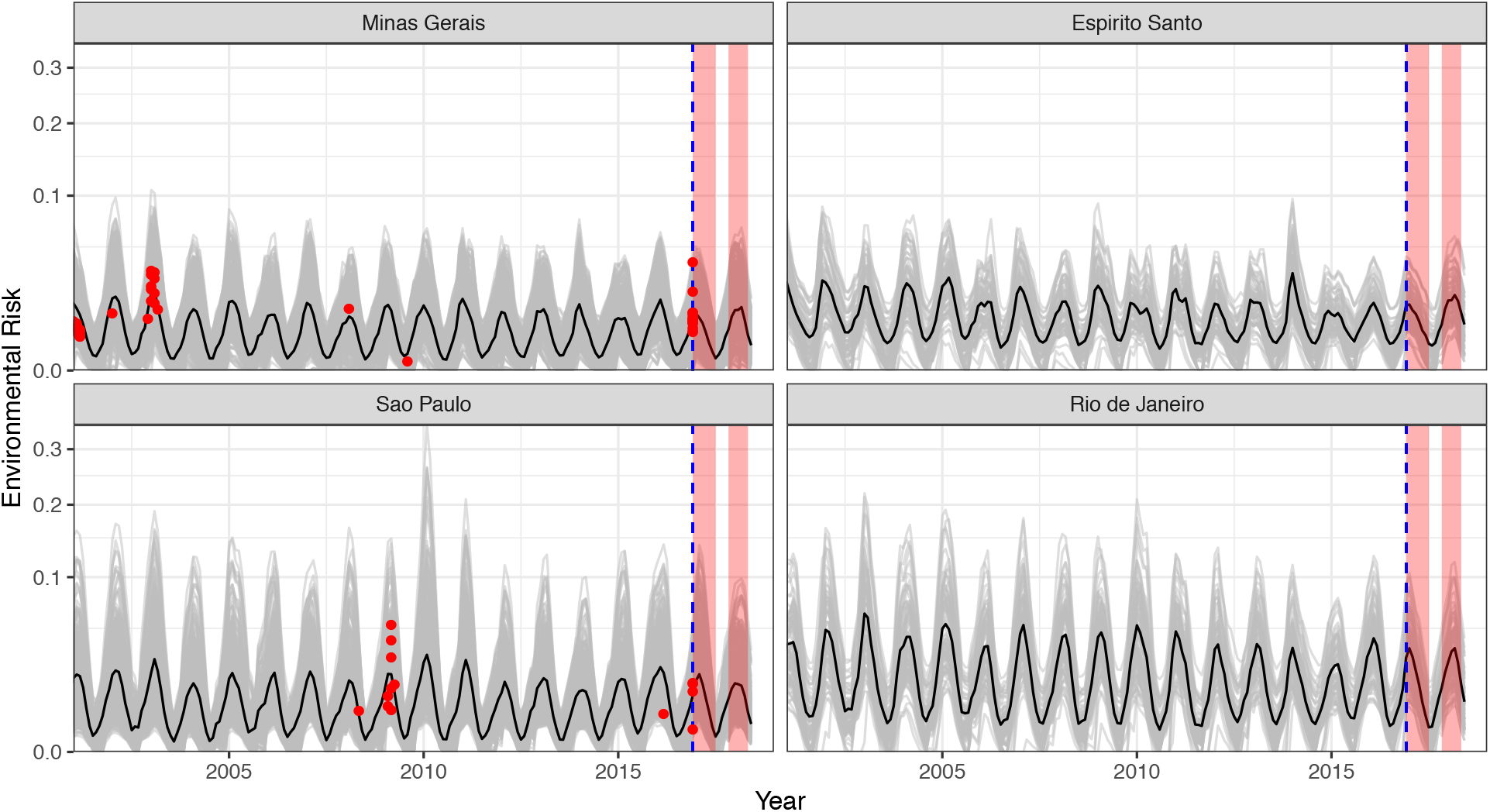
Mechanistic model predicts consistent low, seasonal risk across states in Southeast region of Brazil, where a large outbreak occurred in 2016–2018. Data are only available until the end of 2016 (blue dashed line), so do not include the duration of the 20162018 outbreaks (pink boxes). Only 2001–2016 spillovers are shown, defined as municipality-months with human yellow fever cases (red points). Grey lines are municipality estimates of maximum environmental risk and the black line is the environmental risk averaged over all municipalities in the state. Prior to the large outbreak in 2017–2018, spillover had occurred in Minas Gerais and São Paulo (right panels) but not in Espírito Santo or Rio de Janeiro (left panels).

## Discussion

Our mechanistic understanding of environmental risk of spillover—which combines reservoir host and sylvatic vector distributions, vector contact with reservoirs and humans, and vector dispersal, survival, infectiousness, and seasonal abundance—predicts yellow fever spillover into humans with high accuracy (AUC = 0.72; Fig. 3). Within each municipality and month, the maximum risk, rather than the mean risk, was the best predictor of spillover occurrence, suggesting that local heterogeneity in risk within municipalities is important for determining spillover probability. Rainfall-driven seasonality in the vector populations and temperature-driven seasonality in vector survival and infectiousness accurately predicted seasonal variation in spillover (Fig. 4a). While interannual variation in risk was not well-predicted in the environmental risk model based on climate and land cover information alone, including phenomenologically modeled variation in primate yellow fever infection prevalence improved predictions of year-to-year variation in spillover (AUC = 0.79).

Although we hypothesized that low vaccination coverage and high human population density would each increase spillover risk, neither improved model accuracy for predicting spillover in the mechanistic model (Fig. 3). However, we found that vaccine coverage was the third most important predictor of municipality-months with spillover when allowing for a nonlinear but generally positive relationship between coverage and spillover probability (Fig. 5).

The recent outbreak is also consistent with the ecological processes driving past spillover in this region (Fig. 6). While environmental risk in 2016–2018 was not elevated above historical levels (2001–2015) and spillover had not occurred in in the states of Espírito Santo or Rio de Janeiro during the previous 15 years, it has previously occurred in Minas Gerais and Sao Paulo states in 2001–2003 and 2008–2009. Data from the recent 2016–2018 outbreak past December 2016 are not included in the statistical models because consistent monthly municipality-scale spillover data across the country are not available for that period.

The boosted regression tree analysis, which aimed to detect candidate drivers of spillover that might be missing from our mechanistic model, identified vaccine coverage, current and lagged environmental risk, temperature, population density, and precipitation as important predictors, which together improved upon mechanistic model predictive performance of pathogen spillover (out-of-sample AUC = 0.95). The relative importance of lagged and current environmental risk provides evidence that the mechanistic model captures the potentially nonlinear and interactive relationship between environmental variables that drive spillover in mosquitoes, reservoir hosts, and humans better than the environmental variables alone. One-month lagged environmental risk may be more important than current environmental risk for predicting spillover because of a lag between cases and reporting. Additionally, environmental suitability for reservoir and vectors may drive reservoir infection dynamics, causing a lag between conditions suitable for virus amplification in the primate reservoir and vector populations, and spillover into humans. Moreover, the relative importance of one-month lagged environmental risk creates the potential for forward prediction of spillover. The boosted regression tree also identified municipality-months with spillover to have low temperatures. As mosquito thermal performance traits often have steep drop-offs at high temperatures, temperature variation affects mosquito traits (132). Our mechanistic model using monthly average temperature may overestimate the suitability in warm temperatures and underestimate the suitability in cool temperatures (131), resulting in the decreasing relationship observed between average monthly temperature and spillover in the boosted regression tree.

In a recent publication, Kaul et al. (8) also used a machine learning approach to predict municipality-months with spillover in Brazil and similarly found rainfall and temperature to be important predictors. However, their model also identified primate richness and fire density as important predictors, while our boosted regression tree analysis ranked municipality average primate richness tenth, municipality maximum primate richness fourteenth, one-month lagged fire area ninth, and current fire area twelfth for variable importance out of fourteen variables. Our covariates add to those used by Kaul *et al.* (8) by including vaccine coverage and our mechanistic environmental risk estimate (current and lagged), which boosted regression trees found to be three of the five most important predictors. We expect that our mechanistic environmental risk estimates capture much of the variation attributed to other environmental variables in the Kaul *et al.* model. Despite the differing relative importance of variables for predicting spillover in the two models, they both predict that seasonal patterns vary by regions of Brazil and find Southeast Brazil seasonally suitable for yellow fever spillover. Our mechanistic model further illustrates that this differing seasonality can be explained by seasonal variation in vector survival and infectiousness driven by temperature and vector abundance driven by rainfall.

Given the importance of vaccination campaigns in limiting yellow fever outbreaks, we expected that the number of susceptible (unvaccinated) people would be an important positive predictor of yellow fever spillover occurrence, yet mechanistic population-scaled risk performed worse at predicting spillover than environmental risk alone (Fig. 3). For example, scaling by population size predicts areas of very high risk along the coast of Brazil, where environmental risk is low, but population sizes are high. Additionally, we expected that vaccination coverage and human density might be more predictive of the number of cases in spillover events (for example, the recent outbreak in Southeast Brazil) than the probability of spillover occurring, given that very low environmental suitability will be amplified in large, unvaccinated populations. However, vaccine coverage was not a significant predictor of the number of human cases of yellow fever given that spillover occurred (Supplementary Materials, Table S5). Anecdotally, it is worth noting that prior to the recent large outbreak in Southeastern Brazil in 2016–2018, vaccination rates in the region were low, potentially allowing that outbreak to reach an unusually high magnitude.

The substantial improvement in model prediction from environmental to periodic risk (AUC = 0.72 vs. 0.79) suggests that primate population dynamics, immunity, and infection prevalence may be a key missing component of this mechanistic model. Ongoing surveillance efforts in Brazil are used to detect non-human primate cases of yellow fever as an advanced warning system (133). While this advanced warning system can make a critical difference, the recent outbreaks in Southeast Brazil displayed that in some cases this surveillance may not provide sufficient time to respond to prevent spillover, especially in areas with high populations and low vaccine coverage rates as were found in the Southeast. Incorporating a mechanistic model of non-human primate infection prevalence, driven by local primate surveillance data, could help to indicate when primate cases of yellow fever are likely, to provide additional time for public health officials to respond. This remains a significant and potentially very fruitful gap in our understanding of yellow fever transmission and spillover.

Vector-human contact rates are another important empirical gap in the mechanistic model, which could further refine the relationships between land use, human occupations and behavior, and spillover risk. We approximate human contact rates with sylvatic vectors with percent forest cover, but the relationship is likely much more complex. The surprising decreasing relationship between precipitation and spillover probability in the boosted regression tree (Fig. 5d) may be due to the influence of precipitation on human activities in and around forests, and therefore its influence on human–vector contact (94). Additionally, while vector contacts depend on biting rate of the vector and mosquito biting rates are known to depend on temperature for other species (106,134), we assume constant biting rate in the mechanistic model due to a lack of empirical evidence.

While it was beyond the scope of this paper, the most influential mechanisms in the model could be further identified through sensitivity analyses of specific submodel components. Additionally, associations between model components and spillover probability could be estimated using the framework of percolation models (Washburne *et al.,* this issue). Finally, a thorough uncertainty analysis could highlight the model components most in need of further study to improve prediction of spillover.

Yellow fever is an ancient, historically important human disease that played a central role in the discovery of mosquito transmission of pathogens and the subsequent development of vector control as a public health measure (135). The wealth of existing knowledge about the ecology of yellow fever virus and its sylvatic reservoir hosts and vectors allowed us to synthesize data from 73 published papers to mathematically formalize our ecological understanding of sylvatic transmission and spillover. Although spillover is a stochastic process that is expected to be difficult to predict, the mechanistic model that integrates vector, human host, non-human reservoir, and virus ecology allowed us to predict spillover with surprising accuracy. Historically in the Americas and presently in other regions such as sub-Saharan Africa, yellow fever regularly has entered urban transmission cycles that lead to major human epidemics. The model framework presented here could be extended to include the ecology of different vectors, hosts, and environments, including urban *Ae. aegypti* and more human immune interactions with other flaviviruses, to ask intriguing questions such as: What prevents yellow fever from entering urban transmission cycles in the Americas, where other flavivirus epidemics regularly occur? Why has urban transmission occurred recently in Africa and not in South America? What prevents yellow fever circulation and spillover in Southeast Asia, where sylvatic vectors and non-human primate hosts are present and the climate is suitable? Answers to these questions would further our understanding of the ecology of (re)emerging diseases in different parts of the world. More fundamentally, this work provides clear evidence for the predictive power of mechanistic, ecological models—even for rare events like pathogen spillover—and can provide useful information to enhance public health interventions of zoonotic diseases.

## Supporting information

Risk Estimates

## Additional Information

## Acknowledgements

Freya Shearer kindly provided data on vaccine coverage estimates. We thank the Defense Advanced Research Projects Agency whose funding provided the opportunity to discuss and develop this project at a workshop on pathogen spillover. We would also like to thank the many previous studies who have made their data available as well as publicly available remote sensing data that made this project possible. The manuscript was improved by comments from anonymous reviewers.

## Data Accessibility

All data and code are available on GitHub (https://github.com/marissachilds/YellowFeverSpillover).

## Authors’ Contributions

MLC and EAM conceived of the project and designed the analyses. MLC, NN, and JC collected the data and performed the analyses. NN created the artwork in Fig. 1 and 2. MLC drafted the manuscript. EAM, NN, and JC revised the manuscript and all authors read and approved the final manuscript.

