## Supplementary Materials for "Mosquito and primate ecology predict human risk of yellow fever virus spillover in Brazil"

Figure S4 At each temperature where experiments were performed, we plot observations of mosquito biting that resulted in transmission (1) or no transmission (0) on the y-axis and number of days post infectious blood meal on the x-axis in blue points. The black solid line is the modeled probability of mosquito infectiousness, which takes into account both vector competence and a log-normally distributed EIP. Each panel is labeled by the temperature it represents in degrees Celsius. . . . . . . . . . . . . . . . . . . . . . . . . . . . . . . . . . 12
Figure S5 Each of the parameters govering mosquito infectiousness is modeled as temperature dependent. Vector competence determines the horizontal asymptote of mosquito infectiousness, time to 50% infectious determines the point at which mosquito infectiousness is 50% of the way to vector competence, and standard deviation is the the standard deviation of the exponent of the log-normal distrbution. Solid black lines show model estimates. Blue bars show the range of observed temperatures in lab studies and red bars show range of monthly average temperatures observed in Brazil. . . . . . . . . . . . . . . . . . . . . . . . . . . . . . . . 13

### 1 Spillover model details

#### 1.1 Model details

We approximate environmental risk by discretizing to months rather than continuous time and using a sum over the current month and previous three months:

$$b(\vec{y},t)\beta_h(\vec{y},t)\sum_{\tau=t-3}^{\tau=t}\int_{\vec{x}}\rho_v(\vec{x},\tau)b(\vec{x},\tau)\beta_p(\vec{x},\tau)\kappa(\vec{x},\tau)EIP(T(\vec{x})),t-\tau)s(T(\vec{x})),t-\tau)d(||\vec{y}-\vec{x}||)dxd\tau.$$

Here, $\rho_v(\vec{x},\tau)$ is the density of sylvatic vectors, $b(\vec{x},\tau)$ is the biting rate of vectors, $\beta_p(\vec{x},\tau)$ is the probability of biting a non-human primate, $\kappa(\vec{x},\tau)$ is the non-human primate infection prevalence, $EIP(T(\vec{x})),t-\tau)$ is the probability the vector has completed the extrinsic incubation period and has become infectious, $s(T(\vec{x})),t-\tau)$ is the probability of vector survival, and $d(||\vec{y}-\vec{x}||)$ is vector dispersal. For more detailed variable definitions see Table 1 (Main Text). The model is run in Google Earth Engine (1). The built-in functionality of Google Earth Engine allows for calculations between data sources of differing scales and projections by performing the calculations for a specified output pixel with specified projection and scale. We use the default scale: 1 km x 1 km pixels.

#### 1.2 Data

The data used for the spillover model are described in Table S1.

### 2 Mechanistic sub-model details

#### 2.1 Mosqito density

##### 2.1.1 Methods

We fit species distributions models to combined *Haemagogus janthinomys*, *Hg. leucocelaunus*, and *Sabethes chloropterus* mosquito occurrence data using sampling-bias corrected background points (2). We fit the models using the `maxnet` package in R (3) with a range of regularization parameters (0.5, 1.0, 1.5, 2.0, 2.5, 3.0, 3.5, 4.0, 4.5, 5.0, 6.0, 7.0, 8.0, 9.0, 10.0, 12.5, 15.0, 17.5, 20.0) and feature classes (linear; linear and hinge; linear and quadratic; linear, hinge, and quadratic; linear and product; linear, quadratic, and product; linear, hinge, and product; and linear, hinge, quadratic, and product) and select the model with the lowest small-sample-size corrected Akaike information criterion (AICc).

We use a complementary log-log (cloglog) transform to estimate occurrence probability (4), and calculate mosquito density from occurrence probability ($p$) as $\log(1/(1-p))$ (5).

##### 2.1.2 Data

Covariates are extracted using Google Earth Engine (1). The data used for the species distribution model are described in Table S2.

Occurrence points are from Global Biodiversity Information Facility (6–8) and a search of the literature. We searched in Scopus using the search term "ALL ( ( haemagogus OR sabethes ) AND ( trap* OR collect* OR field OR site OR sample ) )" on June 22, 2018. We searched in Web of Science using the search term "haemagogus OR sabethes" on July 19, 2018. We then limited to papers that caught *Hg. janthinomys*, *Hg. leucocelaunus*, or *Sabethes chloropterus* mosquitoes in South America and reported the GPS location of

Table S1: Data sources for spillover model, including information on the spatial resolution and range, temporal cadence and range, and use of the data.

| **Name** | **Source** | **Spatial Resolution (Spatial Range)** | **Temporal Cadence (Temporal Range)** | **Use** |
|---|---|---|---|---|
| Forest cover | MODIS MOD44B V006 [1] | 250 m (Global) | Yearly (2000-2016) | Used to approximate reservoir-vector and human-vector contact. Assumed 2017 and 2018 identical to 2016 for model estimates of 2017 and 2018. |
| Primate ranges | IUCN [2] | NA (Global) | Static (NA) | Limited to species in *Ateles*, *Aotus*, *Alouatta*, *Saimiri*, *Cebus*, *Callicebus*, *Callithrix*, *Saguinus*, and *Lagothrix* genera [3]. Where no species range maps occurred, reservoir-vector contact rate set to zero. |
| Precipitation | TRMM 3B43 [4] | 0.25 arc degrees (Global) | Monthly (Jan 1998 - Sep 2018) | Used to drive seasonal vector abundance through logistic model fit to field data. NOTE: Used TRMM/3B43V7 image collection available on Google Earth Engine. |
| Human population | CIESIN GPWv4 [5] | 30 arc seconds (Global) | 5 years (2000 - 2020) | Linearly interpolated between 5 year population estimates to determine yearly population estimate. Scales immunological risk to estimate population-scaled risk. NOTE: Used CIESIN/GPWv4/unwpp-adjusted-population-count image collection available on Google Earth Engine. |
| Air temperature | GLDAS-2.1 [6] | 0.25 arc degrees (Global) | 3 hours (Jan 2001 - Oct 2018) | Aggregate to monthly average air temperature. Monthly average air temperature used in estimating vector survival and infectiousness using mechanistic trait models. NOTE: Used NASA/GLDAS/V021/NOAH/G025/T3H image collection available on Google Earth Engine. |
| Vaccine coverage | Freya Shearer (personal communication) | Municipality (South America and Africa) | yearly (2001 - 2016) | Methods for estimating vaccine coverage rates from [7]. We use the coverage estimates from the untargeted, unbiased vaccination scenario and estimate the proportion of the population susceptible to yellow fever as one minus the vaccine coverage. Assumed 2017 and 2018 identical to 2016 for model estimates of 2017 and 2018. |

Table S2: Data sources for species distribution model, including information on the spatial resolution and range, temporal cadence and range, and use of the data.

| **Name** | **Source** | **Spatial Resolution (Spatial Range)** | **Temporal Cadence (Temporal Range)** | **Use** |
|---|---|---|---|---|
| Land surface temperature | MODIS MYD11A1 V006 [8] | 1000 m (global) | 1 day (Mar 2000 - Dec 2018) | Calculated yearly minimum, median, and maximum temperature and for each pixel and averaged over 2001-2017. NOTE: Used MODIS/006/MYD11A1 image collection available on Google Earth Engine. |
| Precipitation | CHIRPS Daily (version 2.0) [9] | 0.05 arc degrees (quasi-global) | 1 day (Jan 1981 - Oct 2018) | Calculated following 3 variables: (1) Yearly total precipitation averaged over 2001-2017. (2) Precipitation of the driest month averaged over 2001-2017. (3) Precipitation of the wettest month averaged over 2001-2017. NOTE: Used UCSB-CHG/CHIRPS/DAILY image collection available on Google Earth Engine. |
| Elevation | NOAA ETOPO1 [10] | 1 arc minute (global) | static (NA) | Used bedrock elevation in meters. NOTE: Used NOAA/NGDC/ETOPO1 image available on Google Earth Engine. |
| Forest Cover | Hansen Global Forest Change v1.5 [11] | 1 arc second (global) | static (NA) | Used percent forest cover from 2000. NOTE: Used UMD/hansen/global_forest_change_2017_v1_5 image available on Google Earth Engine. |
| Land Cover | MODIS MCD12Q1 V006 [12] | 500 meters (global) | yearly (2001 - 2016) | Used FAO-LCCS2 land use layer from 2007. NOTE: Used MODIS/006/MCD12Q1 image collection available on Google Earth Engine. |
| Environmental Vegetation Index | MODIS MOD13A2 V006 [13] | 1000 meters (global) | 16 days (Feb 2000 - Nov 2018) | Calculated median annual EVI averaged over 2001 - 2017. NOTE: Used MODIS/006/MOD13A2 image collection available on Google Earth Engine. |

the capture, resulting in 55 papers (9–63). For the sampling-bias correction, we used Global Biodiversity Information Facility occurrence records from other mosquito species (64), and used locations of other mosquito captures where no occurrence records existed of *Hg. janthinomys*, *Hg. leucocelaunus*, or *Sa. chloropterus* as pseudo-absence points.

###### 2.1.3 Results

The model with the lowest AICc had a regularization parameter of 6.0, and included linear and quadratic features. These were the parameters used to fit the final model. Partial dependence plots showing the marginal response of yellow fever spillover to all covariates are shown in Figure S1. The predicted distribution of yellow fever vectors over all of South America is shown in Figure S2.

##### 2.2 Mosquito seasonality

###### 2.2.1 Methods

For each location and vector species, we calculate the maximum monthly mosquito capture, and relative mosquito capture for each month as the percentage of maximum monthly mosquito capture for that location. Similarly, we calculate relative monthly rainfall. We fit a logistic regression of relative mosquito capture on present and lagged relative rainfall using `glm` in R.

###### 2.2.2 Data

For data on mosquito seasonality, we searched the literature and selected papers with field data of adult mosquito captures in consecutive months that also reported rainfall data. We limited to papers that collected at least one of the three sylvatic yellow fever vectors (*Hg. janthinomys*, *Hg. leucocelaenus*, and *Sa. chloropterus*). We identified 6 papers that fit these criteria (52,57,65–68). When data were reported in graphical form, we use WebPlotDigitizer (69) to extract values.

###### 2.2.3 Results

Results from the logistic regression are shown in Table S3. A comparison of model estimates and data are shown in Figure S3.

Table S3: Coefficients from logistic regression of seasonal relative mosquito abundance on current and lagged relative rainfall.

| | Estimate | Std. Error | Z value | p-value |
|---|---|---|---|---|
| Intercept | -2.565475 | 0.4332215 | -5.921855 | 0.0000000 |
| Lagged relative rainfall | 1.996189 | 0.7288357 | 2.738874 | 0.0061650 |
| Current relative rainfall | 1.582762 | 0.7088461 | 2.232871 | 0.0255574 |

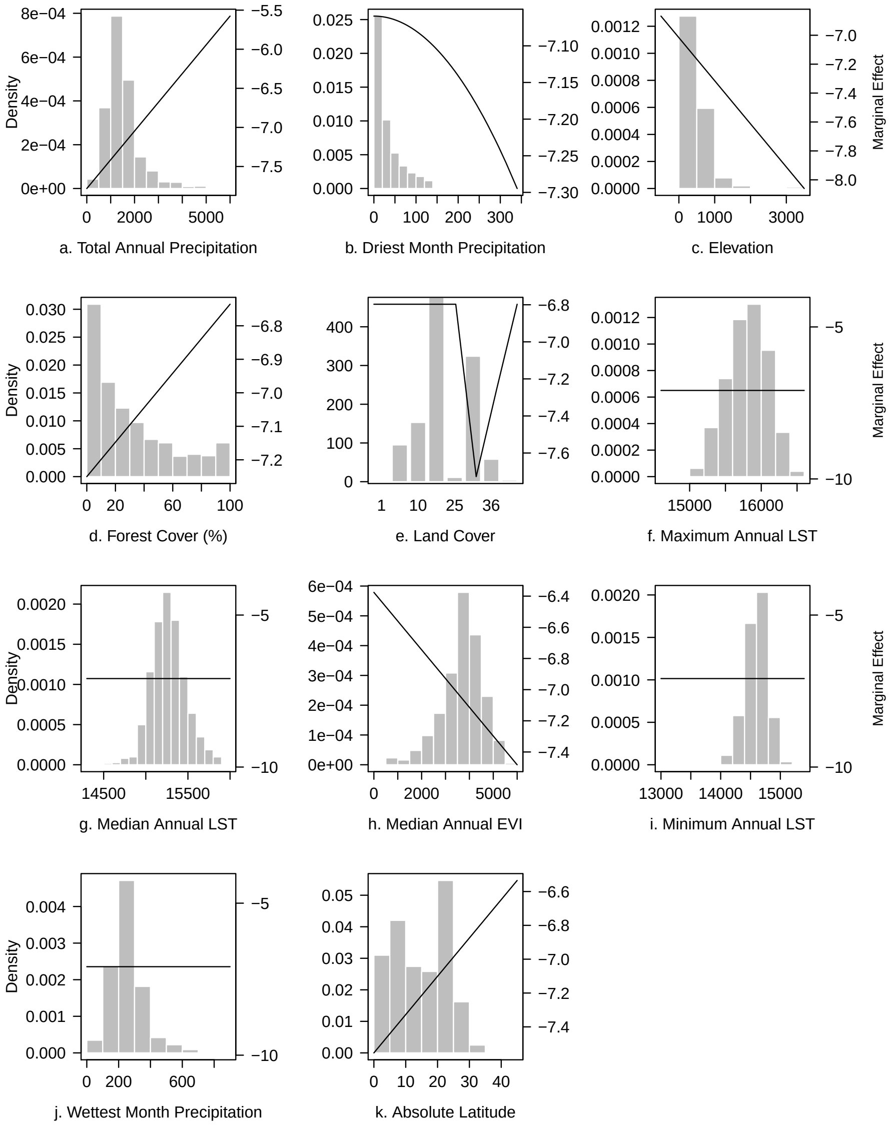

Figure S1: Partial dependence plots of covariates used in species distribution model. Histograms show the distribution of pixels at each covariate value (left axis) and solid lines show the marginal effects of covariate on model prediction (right axis). Covariates with flat marginal effects were identified as unimportant for model prediction. LST Land Surface Temperature, EVI Enhanced Vegetation Index

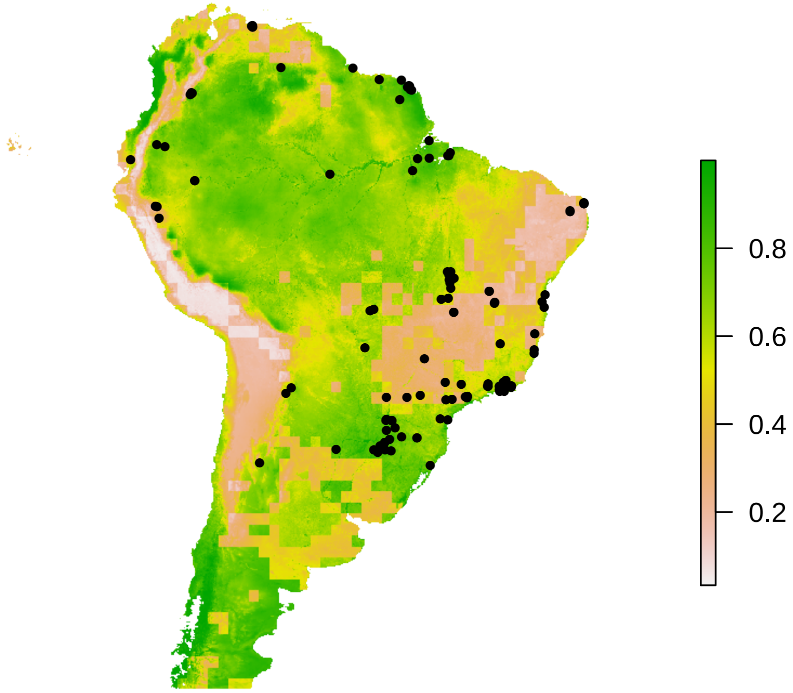

Figure S2: Predicted distribution of vectors from species distribution model, with black points indicating presence locations. Color indicates probability of occurence from highest (green) to lowest (peach).

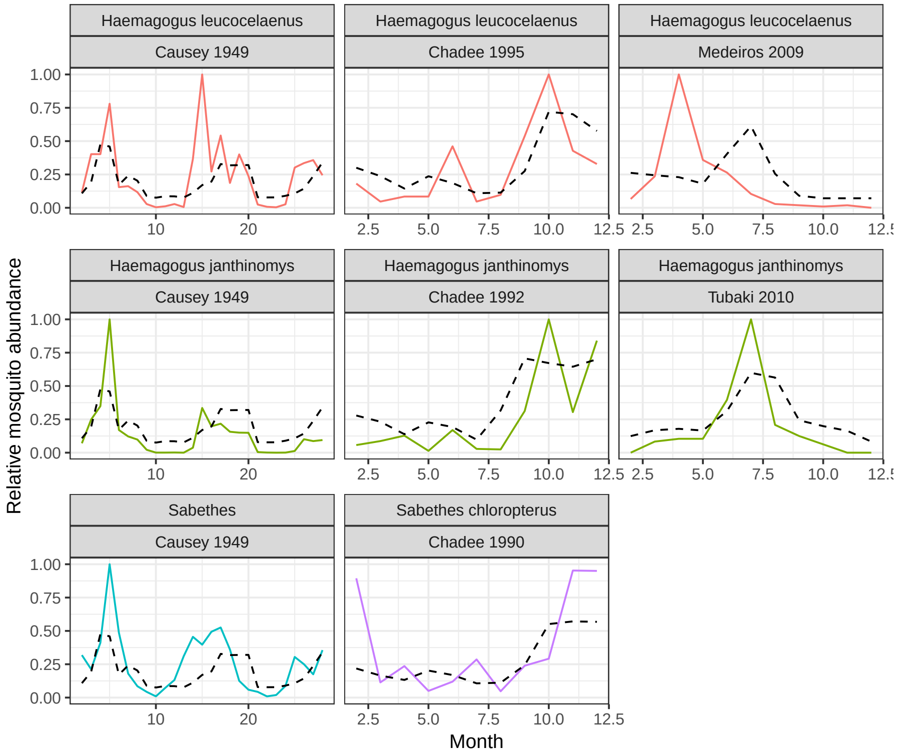

Figure S3: Comparison of mosquito seasonality model to data. Colored lines show data from field studies and dashed black lines show model estimates. Each panel is labeled by the species and study.

#### 2.3 Mosquito survival

##### 2.3.1 Methods

We fit a quadratic function to the relationship between temperature and lifespan (70,71), with differing coefficients for laboratory and field data given the differing lifespans observed in the two settings:

$$L_i \sim N(c_i(T - T_0)(T - T_m), \sigma^2),$$

where $i$ indicates the setting of lab or field. $L_i$ is the lifespan in each setting, $c_i$ is a scaling coefficient, $T_0$ and $T_m$ are the lower and upper critical thermal limits (respectively), and $T$ is the temperature. This assumes that the critical thermal limits are the same in both laboratory and field settings, but that each setting has different maximum lifespan. We use the coefficient from field data for the mechanistic model in the spillover model. The model is fit using the `rstan` package in R (72), and run with 6 chains with 6000 iterations each. We assume that mosquito mortality is constant at a given temperature due to limited information, and calculate daily survival probability as $p = \exp(-1/L)$.

##### 2.3.2 Data

Data are collected from 3 papers (73–75), which were the only sources identified that report *Hg. janthinomys*, *Hg. leucocelaenus*, and *Sa. chloropterus* lifespans and temperatures at which the mosquitoes were reared or caught.

###### 2.3.3 Results

The fitted models for lab and field data are shown in Figure 1e (Main Text).

##### 2.4 Mosquito infectiousness

###### 2.4.1 Methods

Given a set of mosquitoes feed upon an infectious blood meal, we assume that the vector competence, or maximum proportion of mosquitoes becoming infectious, is a quadratic function of temperature (71), and that for each mosquito who becomes infectious, the time to infectiousness has a log-normal probability distribution (76):

$$\begin{aligned} M &= c(T - T_0)(T - T_m) \\ \mu &= \mu_0 + \mu_T T \\ \sigma &= \exp(\sigma_0 + \sigma_T T) \\ EIP &\sim Log - normal(\mu, \sigma), \end{aligned}$$

where $M$ is the maximum proportion infectious, $EIP$ is the time for a mosquito to become infectious, $T$ is the temperature, $T_0$ and $T_m$ are the lower and upper critical thermal limits (respectively), and $c < 0$ is a scaling coefficient. Additionally, $\mu$ is the log of $EIP_{50}$ (time to 50% of max infectious), $\mu_0$ is a scaling factor, and $\mu_T$ is the effect of temperature of $EIP_{50}$. At any point, we model a mosquito's probability of being infectious as $M$ times the cumulative distribution of EIP. The data collected vary in the number of mosquitoes used in each experiment, and mosquitoes were often grouped for biting on primates, so for observations with transmission, we model each censored observation as the probability that at least one mosquito of the group became infectious during the interval and for observations without transmission, we model the censored observation as the probability that none of the mosquitoes became infectious by that time. The model was fit in R using the package `rstan`(72). We run 4 chains with 4000 post-warmup draws per chain and use the median for the parameter estimates.

###### 2.4.2 Data

We collect data (77–83) on yellow fever virus transmission experiments with *Sabethes* and *Haemagogus* species mosquitoes. We use only experiments where mosquito infectivity is tested through bite on a vertebrate. Experimental observations were treated as censored, that is, we either had an interval during which the mosquito or group of mosquitoes became infectious or an interval on which the mosquito or group of mosquitoes did not become infectious during testing (76).

###### 2.4.3 Results

Figure S4 shows transmission experiment data and estimated curves at different temperatures with data. Figure S5 shows vector competence, $EIP_{50}$, and the standard deviation of the log-normal distribution as a function of temperature.

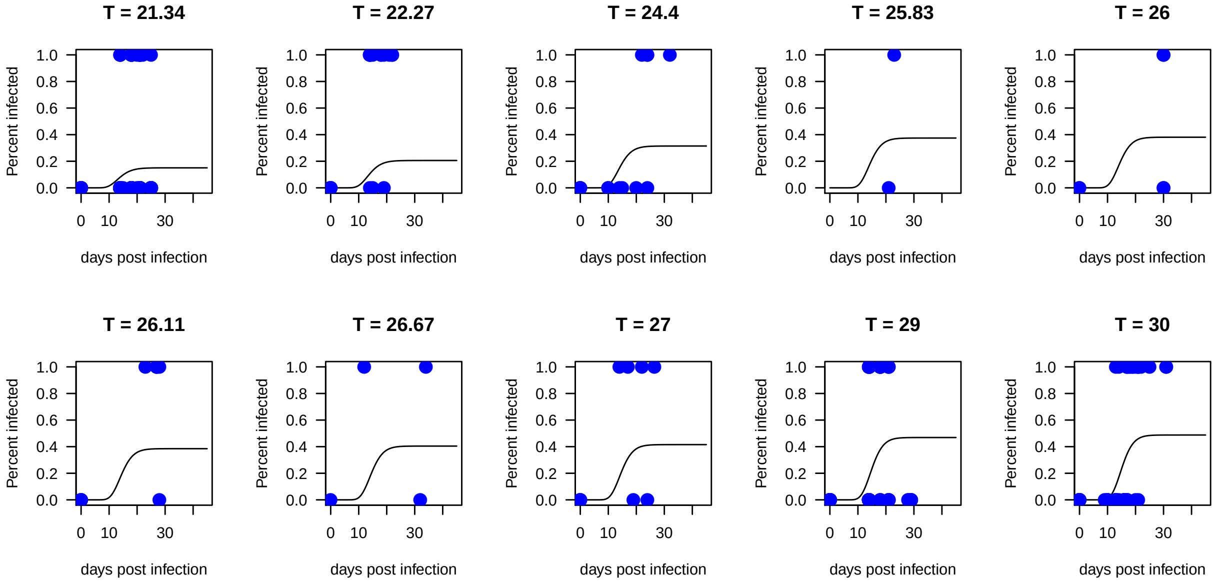

Figure S4: At each temperature where experiments were performed, we plot observations of mosquito biting that resulted in transmission (1) or no transmission (0) on the y-axis and number of days post infectious blood meal on the x-axis in blue points. The black solid line is the modeled probability of mosquito infectiousness, which takes into account both vector competence and a log-normally distributed EIP. Each panel is labeled by the temperature it represents in degrees Celsius.

#### 2.5 Mosquito dispersal

##### 2.5.1 Methods

We use data on mosquito dispersal from a mark-recapture study, and extract capture station locations using WebPlotDigitizer (69). We then fit a negative exponential dispersal kernel (84), with a negative binomial measurement process:

$$Y_i \sim NB\left(T_i\gamma\frac{1}{2\pi\beta}\exp\left(-\frac{r_i}{\beta}\right), k\right),$$

where $Y_i$ is the number of mosquitoes caught at location $i$, $T_i$ is the amount of time spent capturing at location $i$, $r_i$ is the distance of location $i$ from the release location, $\beta > 0$ is the dispersal scaling parameter, $k > 0$ is the overdispersion parameter, and $\gamma > 0$ is a scaling factor to account for both the number of mosquitoes released and the recapture rate. The model is fit using a Bayesian framework in R using the package `rstan`(72). We run 4 chain with 2000 post-warmup draws per chain and use the median for the parameter estimate.

##### 2.5.2 Data

The data used are from a mark-recapture study performed by Causey (85).

##### 2.5.3 Results

Figure S6 compares the mark-recapture dispersal data with the model estimates, and incorporates the time spent in collection at each location.

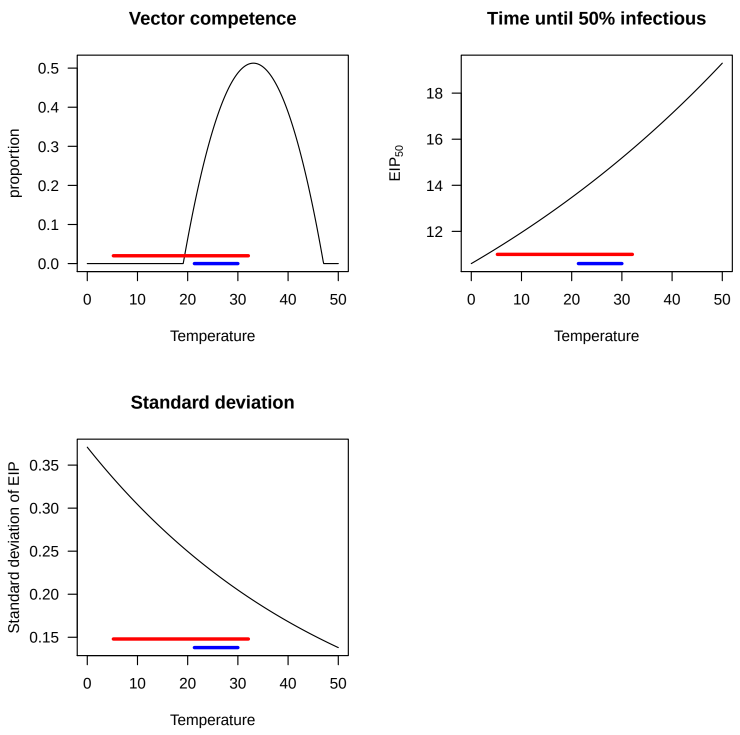

Figure S5: Each of the parameters govering mosquito infectiousness is modeled as temperature dependent. Vector competence determines the horizontal asymptote of mosquito infectiousness, time to 50% infectious determines the point at which mosquito infectiousness is 50% of the way to vector competence, and standard deviation is the the standard deviation of the exponent of the log-normal distrbution. Solid black lines show model estimates. Blue bars show the range of observed temperatures in lab studies and red bars show range of monthly average temperatures observed in Brazil.

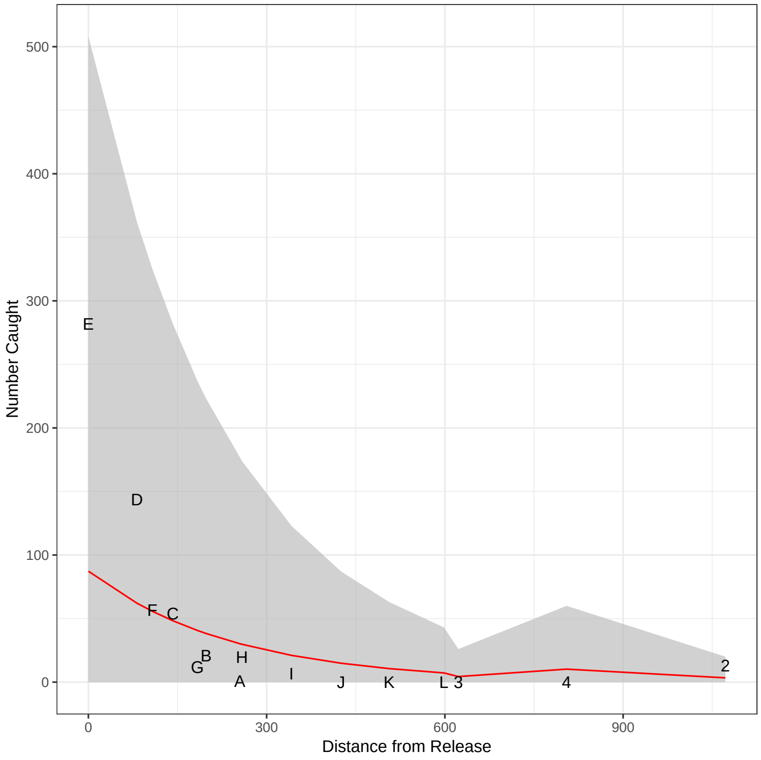

Figure S6: Comparison of dispersal kernel estimate to data. Points are labaled by their trapping location letter from [14]. Red line indicates estimated number of mosquitoes caught at each location from the negative exponential dispersal model when accounting for trapping effort in each sampling location. 95% confidence interval from negative binomial distribution shown in grey shading.

### 3 Phenomenological primate dynamics details

#### 3.1 Methods

We fit a phenomenological sine curve with a seven year period (86) to the yearly number of municipality-months with spillover, and then rescale the curve to be between zero and one, as it represents the reservoir infection prevalence. Using the fact that $A\sin(x+B) = a\sin(x) + b\cos(x)$, we assume that human spillover events are a proxy for infection prevalence during reservoir epizootics and fit a linear model to predict number of municipality-months with spillover each year from the sine and cosine of year, transformed to have a 7 year period.

#### 3.2 Data

We use monthly human cases of yellow fever from the Brazilian Ministry of Health (87). These cases are reported by municipality of infection and month of first symptoms. We consider spillover to have occurred in a municipality-month if at least one case of yellow fever was reported to have originated from that municipality-month. For the purposes of the phenomenological primate dynamics, we sum the number municipality-month reporting spillover for each year.

#### 3.3 Results

The sinusoidal curve explains 52% of the variation in number of municipality-months with spillover (Main text, Figure 2k).

### 4 Model-data comparison details

#### 4.1 Methods

Given the multiple hypotheses, we use the Bonferroni procedure (88) to ensure that the family-wise error rate remains 5% over all 17 hypotheses tested (8 for associations between spillover probability and 9 for associations with number of cases given that spillover occurred.) For each hypothesis tested, we report the adjusted p-value $\min(mp_i, 1)$, where $m$ is the total number of hypotheses and $p_i$ is the p-value from the hypothesis.

#### 4.2 Data

We used the same data of human cases of yellow fever by municipality and month as described above (Phenomenological primate dynamics: Data). We used Brazilian municipality shapefiles from Instituto Brasileiro de Geografia e Estatística (IBGE) (89) for extracting municipality maximum and mean risk metrics.

#### 4.3 Results

See Table S4 for AIC from logistic regression of spillover on risk estimates and AUC from spillover predicted by risk estimates. See Table S5 for results from linear regression of number of reported cases of yellow fever spillover in locations where spillover occurred predicted by model risk estimates and vaccine coverage and Spearman's correlations between risk estimates and number of cases.

Table S4: Model-spillover comparison results. AIC, logistic regression coefficient, logistic regression p-value, and Bonferroni adjusted logistic regression p-value are all reported from a logistic regression of spillover on model estimates and AUC is reported from the receiver operating characteristic curve from predicting spillover with model estimates.

| Risk Metric | Municipality summary | AUC | AIC | Logistic regression coefficient | Logistic regression p-value | Bonferroni adjusted logistic regression p-value |
|---|---|---|---|---|---|---|
| Environmental | mean | 0.705 | 2771.138 | 76.516 | 0.000 | 0.000 |
| Environmental | max | 0.719 | 2735.158 | 18.287 | 0.000 | 0.000 |
| Periodic | mean | 0.776 | 2764.508 | 101.575 | 0.000 | 0.000 |
| Periodic | max | 0.792 | 2731.526 | 22.925 | 0.000 | 0.000 |
| Immunological | mean | 0.597 | 2800.727 | 102.017 | 0.014 | 0.241 |
| Immunological | max | 0.637 | 2786.488 | 27.825 | 0.000 | 0.000 |
| Population-scaled | mean | 0.518 | 2805.175 | -0.049 | 0.704 | 1.000 |
| Population-scaled | max | 0.639 | 2802.825 | 0.003 | 0.019 | 0.322 |

Table S5: Model-case comparison results. Number of reported cases given spillover predicted by risk metrics and vaccine coverage. We also calculate Spearman's rank correlation coefficient for number of cases and risk metrics.

| Risk Metric | Municipality summary | R-squared | Adjusted R-squared | Coefficient | p-value | Bonferroni adjusted p-value | Spearman correlation coefficient |
|---|---|---|---|---|---|---|---|
| Environmental | mean | 0.011 | 0.004 | -43.273 | 0.212 | 1 | 0.008 |
| Environmental | max | 0.020 | 0.012 | -7.408 | 0.098 | 1 | 0.009 |
| Periodic | mean | 0.008 | 0.001 | -47.538 | 0.277 | 1 | 0.011 |
| Periodic | max | 0.015 | 0.008 | -8.076 | 0.145 | 1 | 0.012 |
| Immunological | mean | 0.000 | -0.007 | -23.236 | 0.872 | 1 | 0.004 |
| Immunological | max | 0.004 | -0.003 | -13.085 | 0.465 | 1 | 0.005 |
| Population-scaled | mean | 0.001 | -0.006 | -0.194 | 0.648 | 1 | 0.001 |
| Population-scaled | max | 0.008 | 0.001 | -0.009 | 0.286 | 1 | 0.006 |
| Vaccine Coverage | mean | 0.018 | 0.011 | -2.026 | 0.108 | 1 | 0.010 |

### 5 Boosted regression tree

#### 5.1 Methods

We split the data into training (80%) and test (20%) sets using spatially and temporally balanced sampling with the `BalancedSampling` package in R (90). We fit a boosted regression tree to predict spillover for each municipality-month. We consider tree complexities ranging from 1 to 10, and learning rates of 0.005 and 0.001 and for each pair of parameters identified the number of trees (up to 5000) that minimized cross validation predictive deviance (91). We used the `dismo`, `gbm`, and `pdp` packages for the analysis (92–94).

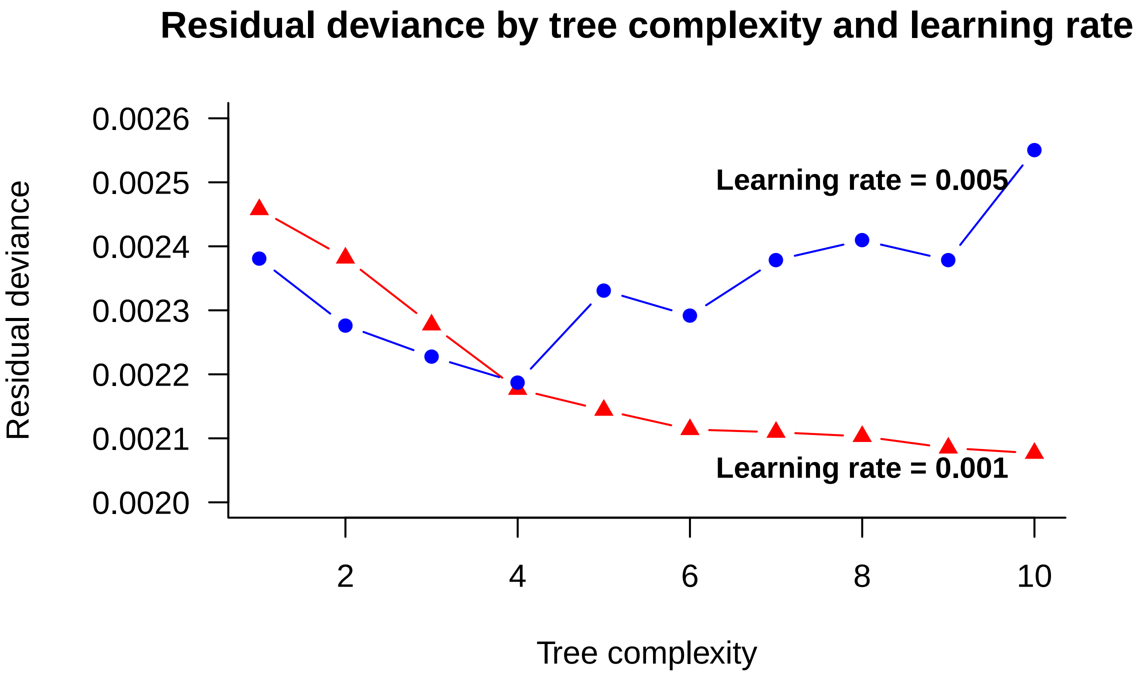

Figure S7: Comparison of predictive deviance across tree complexity and learning rate. Cross validation residual deviance was used to select the optimal number of trees up to 5000 for each set of parameters.

#### 5.2 Data

In addition to lagged and current maximum environmental risk, month of the year, region, vaccine coverage and phenomenological primate dynamics (described previously), we use data on current and lagged fire area, maximum and mean primate species richness in the municipality, population density, fire percent, mean air temperature, and monthly precipitation, as described in Table S6. The same data sources are used for primate species distribution, air temperature, precipitation, and vaccine coverage in both the mechanistic and boosted regression tree models (Table S6 and S1), however the data are used differently due to the differing scales of the models (monthly-pixel for mechanistic and monthly-municipality for boosted regression tree) and differing model forms. For the boosted regression tree, we calculate municipality-month averages of the covariates. We use a different data source for human population distribution in the mechanistic model (CEISIN Gridded Population of the World Version 4, UN-Adjusted Population Count) and boosted regression tree analysis (IBGE municipality population estimates), due to the differing scales required for each model. We capitalized on the non-parametric form of the boosted regression tree to include fire area as an additional covariate that could not be included in the mechanistic model due to limited understanding of the mechanism by which land-use influences spillover risk.

#### 5.3 Results

For comparison of predictive deviance across different tree complexity and learning rate parameters, see Table S7. This comparison can also be seen visually in Figure S7. The set of parameters that minimized cross validation deviance (tree complexity = 10, learning rate = 0.005, number of trees = 5000) was used as the final model. We also show a partial dependence plot for all variables in Figure S8.

Table S6: Data sources for boosted regression tree analysis, including information on the spatial resolution and range, temporal cadence and range, and use of the data.

| **Name** | **Source** | **Spatial Resolution (Spatial Range)** | **Temporal Cadence (Temporal Range)** | **Use** |
|---|---|---|---|---|
| Population Density | IBGE [15] | Municipality (Brazil) | Yearly (2001 - 2016) | We use municipality population estimates and shapefiles of municipalities to determine population density in each municipality. |
| Primate Species Richness | IUCN [2] | NA (global) | static (NA) | Used IUCN species shapefiles to calculate the maximum and spatial average number of primate species in each municipality. Calculations performed in Google Earth Engine [16]. |
| Air temperature | GLDAS-2.1 [6] | 0.25 arc degrees (Global) | 3 hours (Jan 2001 - Oct 2018) | Average temporally then spatially to municipality monthly average air temperature. NOTE: Used NASA/GLDAS/V021/NOAH/G025/T3H image collection available on Google Earth Engine and performed calculation in Google Earth Engine [16]. |
| Precipitation | TRMM 3B43 [4] | 0.25 arc degrees (Global) | Monthly (Jan 1998 - Sep 2018) | Averaged spatially to get municipality average precipitation. NOTE: Used TRMM/3B43V7 image collection available on Google Earth Engine and performed calculation in Google Earth Engine [16]. |
| Fire Area | MODIS MCD64A1 V006 [17] | 500 meters (Global) | Monthly (Nov 2000 - Oct 2018) | We calculate total area of pixels identified as burned for each month. We also calculate percent fire area by dividing by municipality area. NOTE: Used MODIS/006/MCD64A1 image collection available on Google Earth Engine and performed calculation in Google Earth Engine [16]. |
| Vaccine coverage | Freya Shearer (personal communication) | Municipality (South America and Africa) | yearly (2001 - 2016) | Methods for estimating vaccine coverage rates from [7]. We use the coverage estimates from the untargeted, unbiased vaccination scenario and estimate the proportion of the population susceptible to yellow fever as one minus the vaccine coverage. |

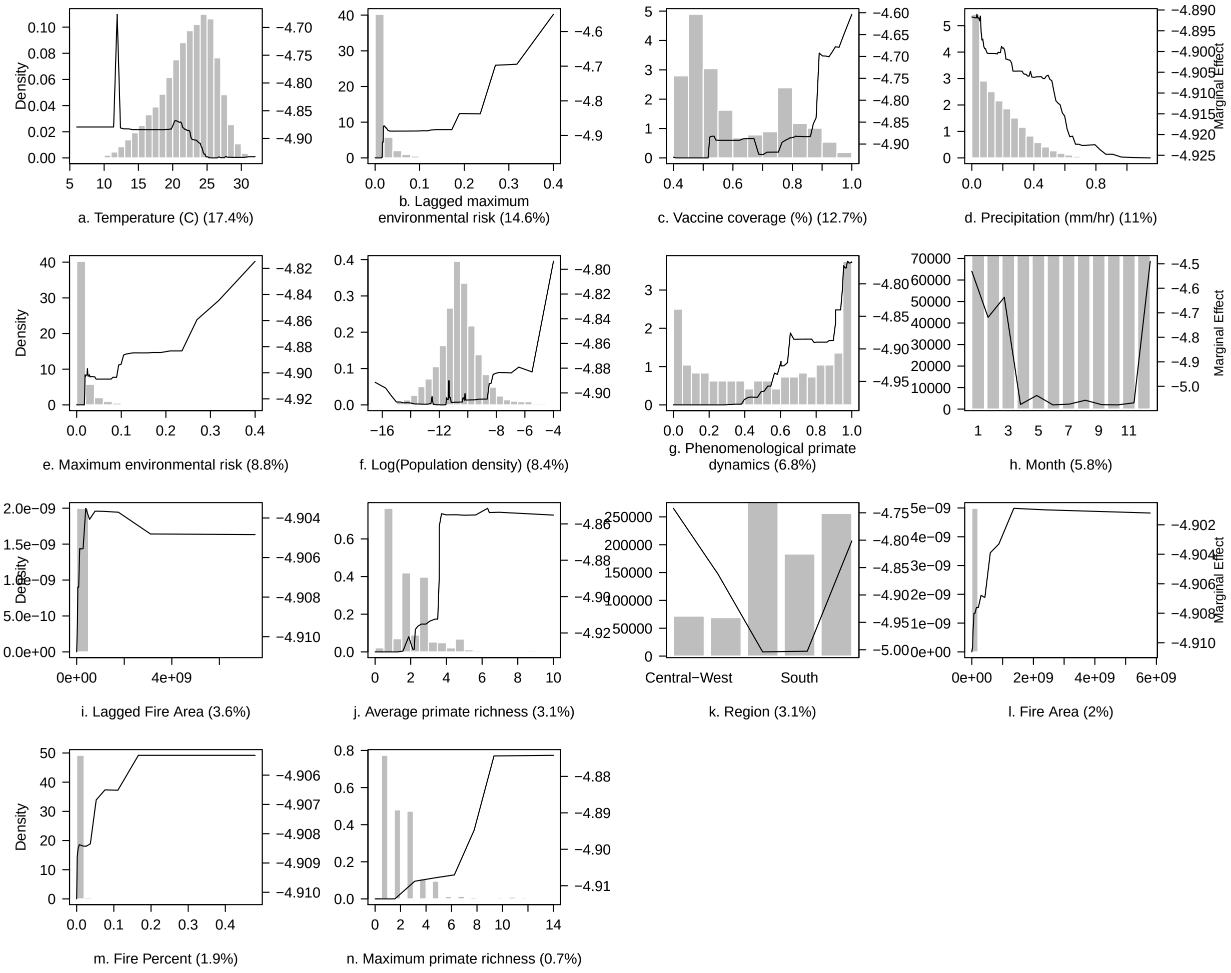

Figure S8: Partial dependence plots of all variables included in the boosted regression tree analysis with histograms showing the distribution of each covariate (left axis) and solid black line showing marginal effect of covariate on model prediction of spillover (right axis).

Table S7: Comparison of predictive deviance across boosted regression tree parameters.

| Tree complexity | Learning rate | Number of trees | Predictive deviance |
|---|---|---|---|
| 10 | 0.001 | 5000 | 0.0020767 |
| 9 | 0.001 | 5000 | 0.0020850 |
| 8 | 0.001 | 5000 | 0.0021030 |
| 7 | 0.001 | 5000 | 0.0021095 |
| 6 | 0.001 | 5000 | 0.0021138 |
| 5 | 0.001 | 5000 | 0.0021440 |
| 4 | 0.001 | 5000 | 0.0021767 |
| 4 | 0.005 | 4600 | 0.0021869 |
| 3 | 0.005 | 5000 | 0.0022276 |
| 2 | 0.005 | 5000 | 0.0022764 |
| 3 | 0.001 | 5000 | 0.0022773 |
| 6 | 0.005 | 1500 | 0.0022916 |
| 5 | 0.005 | 900 | 0.0023311 |
| 7 | 0.005 | 900 | 0.0023785 |
| 9 | 0.005 | 900 | 0.0023785 |
| 1 | 0.005 | 5000 | 0.0023805 |
| 2 | 0.001 | 5000 | 0.0023818 |
| 8 | 0.005 | 900 | 0.0024094 |
| 1 | 0.001 | 5000 | 0.0024575 |
| 10 | 0.005 | 1050 | 0.0025501 |

11. Pecor JE, Jones J, Turell MJ, Fernandez R, Carbajal F, O'Guinn M, et al. Annotated checklist of the mosquito species encountered during arboviral studies in Iquitos, Peru (Diptera: Culicidae). Journal of the

American Mosquito Control Association. 2000;16(3):210–8.

26. Cardoso JDC, Paula MB de, Fernandes A, Santos E dos, Almeida MAB de, Fonseca DF da, et al. Ecological aspects of mosquitoes (Diptera: Culicidae) in an Atlantic forest area on the north coast of Rio

Grande do Sul State, Brazil. Journal of Vector Ecology. 2011;36(1):175–86. https://doi.org/10.1111/j.1948-7134.2011.00155.x

40. Souza RP de, Petrella S, Coimbra TLM, Maeda AY, Rocco IM, Bisordi I, et al. Isolation of yellow fever virus (YFV) from naturally infected haemagogus (conopostegus) leucocelaenus (diptera, culicidae) in

São Paulo state, Brazil, 2009. Revista do Instituto de Medicina Tropical de Sao Paulo. 2011;53(3):133–9. https://doi.org/10.1590/S0036-46652011000300004

55. Pauvolid-Corrêa A, Tavares FN, Alencar J, Silva JDS, Murta M, Serra-Freire NM, et al. Preliminary investigation of culicidae species in South Pantanal, Brazil and their potential importance in arbovirus

transmission. Revista do Instituto de Medicina Tropical de Sao Paulo. 2010;52(1):17–23. https://doi.org/10.1590/S0036-46652010000100004

71. Mordecai EA, Cohen JM, Evans MV, Gudapati P, Johnson LR, Lippi CA, et al. Detecting the impact of temperature on transmission of Zika, dengue, and chikungunya using mechanistic models. Althouse B, editor.

PLOS Neglected Tropical Diseases [Internet]. 2017 Apr;11(4):e0005568. https://doi.org/10.1371/journal.pntd.0005568

86. Camara FP, Gomes ALBB, Carvalho LMF de, Castello LGV. Dynamic behavior of sylvatic yellow fever

in Brazil (1954-2008). Revista da Sociedade Brasileira de Medicina Tropical [Internet]. 2011 Jun;44(3):297–9. https://doi.org/10.1590/S0037-86822011005000024
