## Supplementary figures and images for "Mosquito and primate ecology predict human risk of yellow fever virus spillover in Brazil"

### Risk Estimates

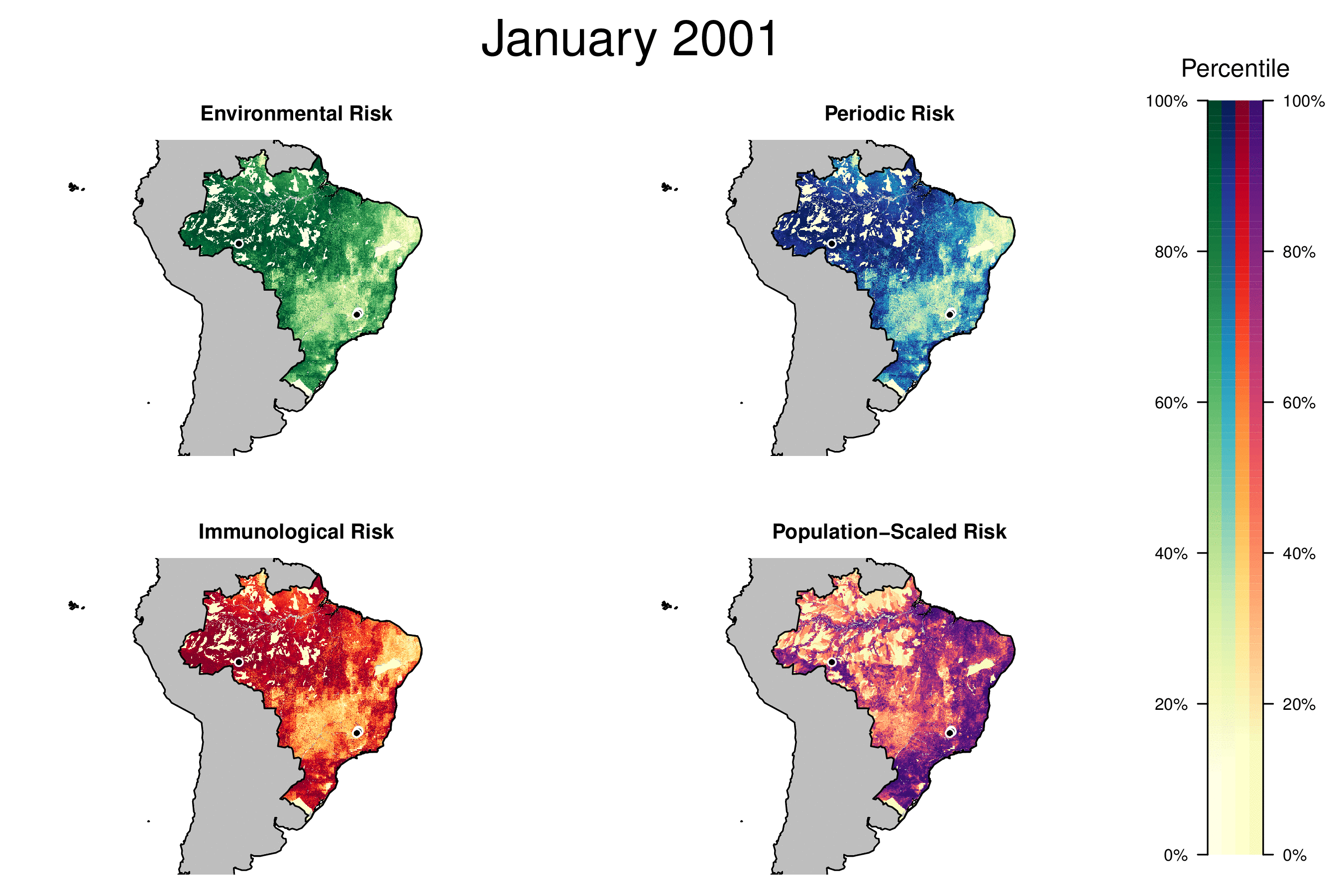
